# Brain dynamics exhibit scale-dependent reversibility during consciousness transitions

**DOI:** 10.64898/2026.08.12.744449

**Authors:** Xu Han, Xiaoai Chen, Samuel R. Cramer, Yizhuo Ding, Nanyin Zhang

## Abstract

Consciousness is a dynamic brain state, yet the systems-level mechanisms underlying transitions into and out of unconsciousness remain poorly understood. It is unclear whether neural dynamics during loss of consciousness (LOC) and recovery of consciousness (ROC) simply retrace the same trajectory or instead follow distinct paths across multiple spatial scales. Here, we simultaneously measured local electrophysiology, whole-brain functional MRI, and pupil dynamics in rats during graded propofol anesthesia to characterize consciousness transitions from local circuits to whole-brain networks. We found that local field potential, regional BOLD responses, and pairwise functional connectivity exhibited largely reversible changes between LOC and ROC. In contrast, the global brain organization showed distinct and asymmetric patterns during the two transitions, as consistently revealed by traveling-wave propagation, low-dimensional network trajectories, and graph-theoretical analyses. Importantly, brain-wide coupling between pupil dynamics and regional BOLD activity remained highly consistent during LOC and ROC, indicating that these distinct global trajectories cannot be simply explained by differences in neuromodulatory tone. Together, our findings identify scale-dependent reversibility as a systems-level organizing principle of consciousness transitions. These results suggest that recovery of consciousness is an active process of large-scale network reorganization rather than merely the reversal of anesthetic suppression.

## Main

Consciousness enables an organism to maintain awareness of itself and the environment. Although the exact neural mechanism remains incompletely understood, it is generally believed that consciousness change is a dynamical brain state. Transitions of the loss of consciousness (LOC) and recovery of consciousness (ROC) both involve coordinated activity across distributed brain systems ^1–3^. As a result, elucidating the spatiotemporal dynamics of brain network activity during the moments into and out of unconsciousness represents a critical step to deciphering the systems-level mechanism underlying consciousness.

Anesthesia-induced unconsciousness (AIU) provides a reversible, conveniently controlled experimental framework to study consciousness. A large body of previous studies has demonstrated that AIU is accompanied by profound neural changes at regional, circuitry and network levels. Electrophysiological studies consistently demonstrate increased low-frequency oscillatory power in local field potentials (LFPs)^4^, along with enhanced cortical synchronization^5–8^ during unconscious states. Neuroimaging studies have revealed widespread changes in brain activity and functional connectivity in thalamocortical, fronto-parietal and default-mode networks^9–13^. Importantly, most of these findings are consistent across anesthetic agents and species, suggesting they represent a general pattern of AIU, as opposed to agent-or species-specific phenomena. Although these findings have substantially advanced our understanding of the neural correlates of unconsciousness, most of these previous studies focused on steady-state changes between wakefulness and AIU. Such approaches provide limited insight into the dynamic processes when consciousness is lost and regained.

A central unresolved question is whether the dynamic processes underlying the LOC and ROC follow simple time-reversed symmetric trajectories or instead exhibit hysteresis and direction-dependent brain organization (Fig. 1A). This distinction is fundamental, as it determines whether consciousness transitions can be explained primarily by graded pharmacological suppression or instead are characterized by an active process of history-dependent brain network reconfiguration. Electrophysiological data show that propofol-induced LOC is marked by a progressive emergence of frontal alpha waves, increased delta-band power, and phase–amplitude coupling, whereas ROC involves a asymmetric dissolution of these LFP structures, including transient intermediate states and altered coupling dynamics, even at comparable anesthetic concentrations^5,14^. Behavioral data also reveal hysteresis in anesthetic responsiveness, evidenced by distinct temporal profiles between LOC and ROC^14,15^. Meanwhile, several neural signatures of anesthesia at the local level, such as changes in LFP power and regional brain activity, exhibit largely reversible patterns between LOC and ROC^9,16^. Taken together, these findings raise a critical systems-level question: although local neural and regional signals may recover symmetrically, does the global spatiotemporal organization of brain activity follow the same trajectory during LOC and ROC (Fig. 1A)?

**Figure 1.**
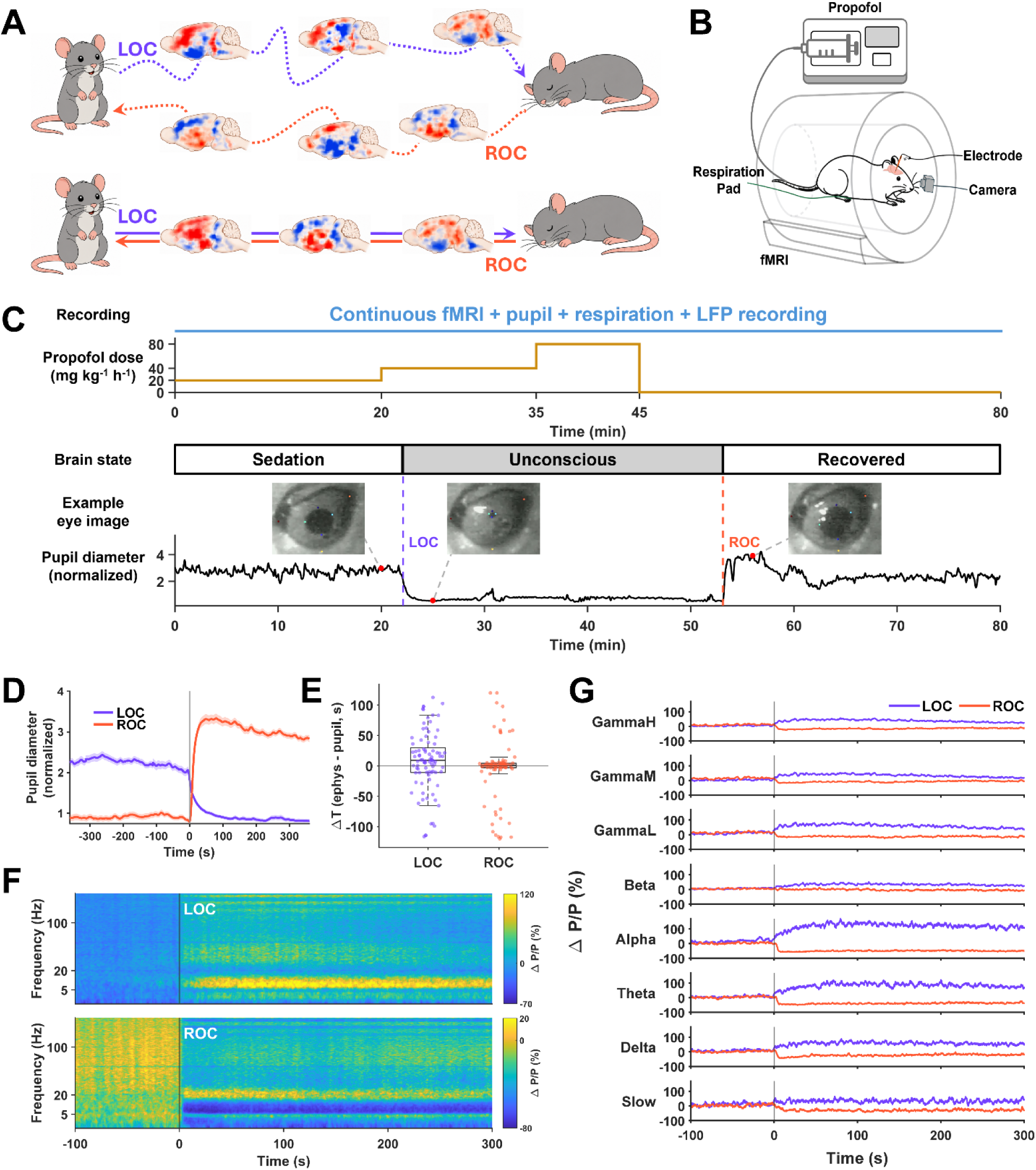
Pupil size changes coincide with established electrophysiological signatures of anesthesia-induced unconsciousness. **(A)** Alternative hypotheses for brain-state transitions during loss and recovery of consciousness. The upper schematic illustrates a transition-dependent hysteretic model, while the lower schematic illustrates a reversible model. **(B)** Schematic illustration of the experimental setup showing a rat in the MRI scanner with a linear electrode implanted in the M2 cortex and dorsal hippocampus, a respiration pad under the chest for breath monitoring, and an infrared camera for pupil tracking. **(C)** Experimental paradigm. Propofol was infused in a stepwise manner at 20 mg kg⁻¹ h⁻¹ for 20 min, 40 mg kg⁻¹ h⁻¹ for 15 min, and 80 mg kg⁻¹ h⁻¹ for 10 min, followed by cessation of infusion and 35 min of recovery. fMRI data, pupil video, respiration signal, and LFP recordings were simultaneously acquired throughout the 80-min session. A representative pupil trace is shown below the dosing timeline. LOC and ROC are marked by vertical lines and were defined based on abrupt changes in pupil diameter. **(D)** Averaged pupil diameter time courses aligned to pupil-defined LOC and ROC transition points. Shaded regions represent mean ± SEM across trials. **(E)** Comparison of pupil-defined transition timing with independently estimated alpha-band LFP transitions in M2. Group distributions are summarized with box, and each dot represents one scan session. Positive values indicate that the LFP transition occurred after the pupil transition, whereas negative values indicate that it occurred before the pupil-defined transition. **(F)** Averaged M2 cortical spectrograms aligned to pupil-defined LOC and ROC. Spectral power is shown as percent change relative to the pre-transition baseline. LOC is associated with a broad increase in LFP power across frequency bands, whereas ROC shows an opposite transition pattern. **(G)** Band-limited M2 LFP power changes aligned to pupil-defined LOC and ROC for canonical frequency bands (Slow, Delta, Theta, Alpha, Beta, GammaL, GammaM, GammaH). Curves summarize the mean LFP power across scans, with shaded bands indicating SEM. Peak post-transition ΔP/P values for LOC were 60.5%, 74.6%, 107.3%, 147.7%, 44.5%, 78.5%, 49.8%, and 53.0%, respectively; corresponding values for ROC were −39.0%, −42.0%, −48.3%, −57.0%, −10.1%, −21.6%, −16.6%, and −22.8%.

A technical challenge to systematically investigate the dynamic processes of LOC and ROC has been the lack of simultaneous, multimodal measurements capable of capturing local neural activity, whole-brain dynamics, and arousal-related physiological signals during rapid state transitions. Local electrophysiological recordings provide direct measures of neuronal population activity and oscillatory dynamics with high temporal resolution but are spatially limited. Functional MRI (fMRI), on the other hand, has brain-wide coverage with high spatial resolution and can measure large-scale brain network activity, yet has slow temporal resolutions. Moreover, autonomic processes play a critical role in regulating arousal and consciousness, but are often inferred indirectly or omitted. Therefore, integrating these complementary modalities represents a powerful framework for a comprehensive understanding of consciousness transitions.

Here, we address this challenge by combining simultaneous electrophysiology, whole-brain fMRI, and pupillometry in a rat model undergoing graded propofol anesthesia. By leveraging the established capability of simultaneous electrophysiology-fMRI-physiological signal measurement^4,17,18^, we recorded LFPs from the hippocampus and cortex (secondary motor cortex, M2) while acquiring concurrent whole-brain fMRI and pupil measurements as animals transitioned from wakefulness to unconsciousness through graded anesthetic administration and subsequently recovered following anesthetic discontinuation. Pupil size was measured as it provides a sensitive, noninvasive index of arousal and neuromodulatory tone^19^. This experimental approach enabled precise temporal alignment of local neural activity, global brain dynamics, and autonomic arousal across both LOC and ROC. We found that pupil size closely tracks transitions into and out of unconsciousness and aligns with robust changes in local neural activity, as reflected by increases in LFP power during LOC and decreases during ROC. We then examined how brain-wide activity and functional connectivity evolved during these transitions and tested the hypothesis that LOC and ROC would be associated with distinct spatiotemporal patterns of network reconfiguration, reflecting direction-dependent state transitions rather than simple time-reversed dynamics.

## Results

To systematically characterize the spatiotemporal dynamics of brain-wide activity during LOC and ROC, we simultaneously recorded local electrophysiology, whole-brain fMRI, and autonomic responses in rats undergoing graded propofol anesthesia (Fig. 1B,C; 12 rats, 90 recording sessions). Following a 20 mg kg⁻¹ propofol bolus administered 30 min before imaging, propofol was administered in stepwise increments (low dose: 20 mg kg⁻¹ h⁻¹ for 20 min; medium dose: 40 mg kg⁻¹ h⁻¹ for 15 min; high dose: 80 mg kg⁻¹ h⁻¹ for 10 min), followed by cessation of infusion and a 35-min recovery period (Fig. 1C). Using a similar anesthesia protocol, we previously demonstrated that LOC, defined by loss of righting reflex and electrophysiology recording, occurs during the medium-dose infusion (40 mg kg⁻¹ h⁻¹)^4^. Accordingly, animals remained behaviorally awake during the low dose and recovered consciousness following cessation of propofol administration (Fig. 1C).

Electrophysiological signals from the secondary motor cortex (M2) and dorsal hippocampus were continuously recorded alongside fMRI, pupillometry, and respiration throughout the experiment (Fig. 1B,C). MRI-related artifacts in the raw electrophysiological recordings were removed using a template regression approach previously established in our laboratory^4,17,20^, yielding high-quality denoised signals suitable for spectral analysis. Pupil landmarks were tracked using DeepLabCut^21^ (Fig. 1C,D), and the pupil diameter was computed from the tracked coordinates using an in-house MATLAB pipeline.

### Pupil size closely tracks transitions into and out of unconsciousness

To precisely identify the moments of LOC and ROC, we combined pupil dynamics with simultaneous local electrophysiological recordings. Because pupil size provides a noninvasive index of arousal and is closely associated with brainstem neuromodulatory activity ^22–24^, we examined whether changes in pupil diameter coincided with established electrophysiological signatures of anesthesia-induced unconsciousness.

Representative pupil images acquired during sedation, unconsciousness, and recovery illustrate marked pupil constriction during LOC and re-expansion during recovery (Fig. 1C, D), indicating that pupil diameter is highly sensitive to consciousness state transitions. Using these transition-related pupil dynamics, LOC was defined as the prominent negative peak in the first derivative of pupil diameter at the onset of sustained pupil constriction, whereas ROC was defined analogously as the prominent positive peak at the onset of sustained pupil dilation. Quantitatively, mean pupil diameter (Fig. 1D) decreased by 62.6% during LOC (95% CI, 60.2–65.0%) and increased by 493.0% during ROC (95% CI, 437.0– 549.0%).

To assess whether pupil-defined transitions corresponded to neural state transitions, we compared their timings with independently estimated LFP transitions in the secondary motor cortex and dorsal hippocampus. LFP-defined LOC and ROC were identified as the times of the steepest increase and decrease, respectively, in alpha-band power, which are well-established electrophysiological signatures of anesthetic-induced consciousness state transitions^4,5,25^. In M2, the mean signed timing difference (LFP-defined time minus pupil-defined time) was 7.8 s for LOC (95% CI, −2.6 to 18.2 s) and −4.4 s for ROC (95% CI, −14.5 to 5.7 s; Fig. 1D), indicating no systematic temporal offset between the two measures (Fig. 1E). A similar temporal correspondence was observed for LFP-defined transitions in the dorsal hippocampus (Fig. S1A). Respiration rate also showed abrupt changes around the pupil-defined LOC and ROC time points (Fig. S1B), providing additional physiological support for these transition markers. We therefore used pupil-defined transition times as the common temporal reference for all subsequent multimodal analyses.

Using pupil-defined LOC, power spectral analysis revealed broad-band increases in LFP power across all canonical frequency bands (slow wave [SW], delta, theta, alpha, beta, low gamma [GammaL], middle gamma [GammaM], and high gamma [GammaH]). In both cortical (Fig. 1F, G) and hippocampal (Fig. S1C, D) recordings, theta-and alpha-band activity exhibited the largest relative increases compared with other frequency bands (peak post-LOC ΔP/P in M2: SW 60.5%, Delta 74.6%, Theta 107.3%, Alpha 147.7%, Beta 44.5%, GammaL 78.5%, GammaM 49.8%, GammaH 53.0%; in hippocampus: SW 19.2%, Delta 50.1%, Theta 74.8%, Alpha 106.0%, Beta 39.1%, GammaL 36.8%, GammaM 20.5%, GammaH 15.7%).

Mirroring the coupling observed during LOC, pupil-defined ROC displayed broad-band reduction in LFP power. In the cortex, peak reductions in band-limited power were observed across all frequency bands (SW −39.0%, Delta −42.0%, Theta −48.3%, Alpha −57.0%, Beta −10.1%, GammaL −21.6%, GammaM −16.6%, GammaH −22.8%; Fig. 1F, G). Similar but smaller reductions were observed in the hippocampus (SW −21.4%, Delta −29.7%, Theta −37.0%, Alpha −46.1%, Beta −7.0%, GammaL −11.8%, GammaM −5.1%, GammaH −6.7%; Fig. S1C, D).

Together, these findings demonstrate that pupil-defined transitions are tightly coupled to rapid changes in neural spectral activity across cortical and hippocampal regions. This cross-modal temporal correspondence supports the use of pupil dynamics as a reliable physiological marker for aligning subsequent analyses of LOC and ROC. Moreover, the broadly reversed spectral responses across the two transitions indicate substantial reversibility of local neural activity during loss and recovery of consciousness.

### Brain-wide fMRI activity is organized into a limited set of spatiotemporal motifs during LOC and ROC

Using pupillometry-defined transition points, we next characterized whole-brain BOLD dynamics surrounding LOC and ROC (Supplementary Movies 1 and 2). To facilitate systems-level analysis, the brain was parcellated into 77 regions of interest (ROIs) grouped into 11 anatomical systems.

Whole-brain BOLD activity exhibited highly structured yet heterogeneous temporal responses around both LOC and ROC as shown in Fig. 2A. Temporal profiles of individual ROIs within each anatomical system are also plotted in Fig. S2.

**Figure 2.**
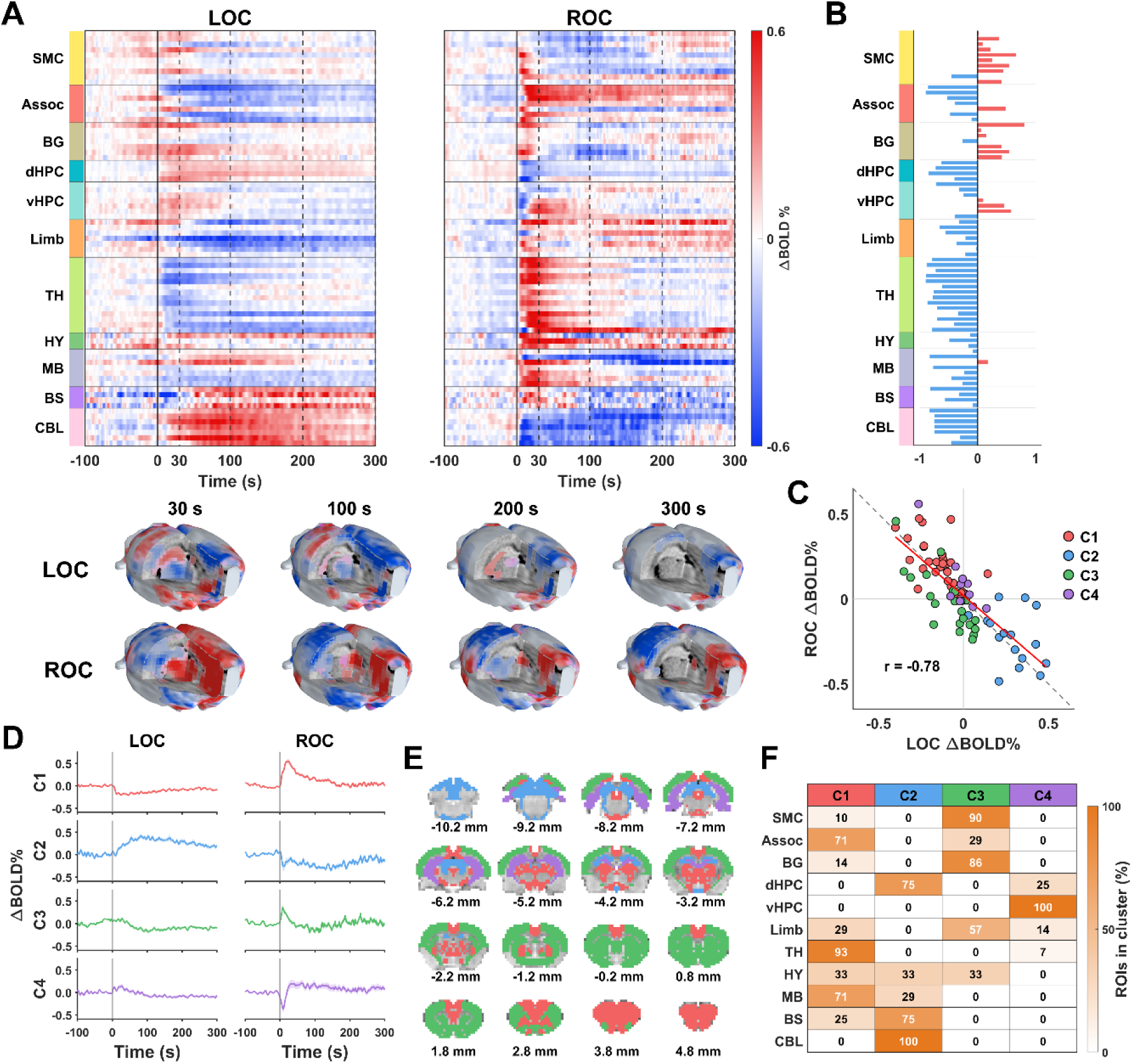
ROI-wise BOLD dynamics reveal reversible but spatially heterogeneous transition motifs during LOC and ROC. **(A)** ROI-wise BOLD signal changes aligned to LOC and ROC transition points. Each row represents the BOLD time course of one ROI, grouped by anatomical system. BOLD signals are shown as the percent change relative to the pre-transition baseline. Representative 3D brain maps below the heatmaps show the spatial distribution of BOLD changes at selected time points (30, 100, 200, and 300 s) after LOC or ROC. Abbreviation: SMC, sensorimotor cortex; Assoc, polymodal association cortex; BG, basal ganglia; dHPC, dorsal hippocampus; vHPC, ventral hippocampal–parahippocampal system; Limb, non-hippocampal limbic system; TH, thalamus; HY, hypothalamus; MB, midbrain; BS, brainstem; CBL, cerebellum. **(B)** Reversibility of ROI-wise BOLD responses. Each horizontal bar represents the correlation between LOC-and ROC-aligned BOLD time courses for one ROI. Negative values indicate opposite LOC and ROC response patterns, whereas positive values indicate similar response patterns. **(C)** Relationship between LOC and ROC response amplitudes across ROIs. Each dot represents one ROI and is colored by the K-means clustering membership. The gray dashed line indicates the y=−x reference line, representing perfect reversal with equal magnitude between LOC and ROC. **(D)** Averaged BOLD time courses for four ROI clusters identified from transition-related dynamics. For each cluster, mean ± SEM LOC-and ROC-aligned time courses are displayed side by side. **(E)** Spatial distribution of ROI cluster membership overlaid on coronal brain slices, illustrating the anatomical organization of distinct BOLD response motifs. **(F)** Percentage of ROIs from each anatomical system assigned to each cluster.

Although individual brain regions displayed distinct activity profiles (Fig. 2A), many regions exhibited strikingly reversed temporal dynamics between the two transitions. To quantitatively assess the reversibility, we calculated the temporal correlation between LOC-and ROC-aligned BOLD responses for every ROI (Fig. 2B). Most brain systems exhibited strong negative correlations, indicating highly reversible temporal activity patterns between LOC and ROC. The thalamus, polymodal association cortex, dorsal hippocampus, and cerebellum showed the strongest negative correlations, and this temporal reversibility was highly consistent across individual ROIs within each system. In contrast, a minor portion of brain systems exhibited direction-dependent responses, reflected by relatively weak temporal correlations between LOC and ROC.

Spatial BOLD patterns were also broadly opposite in polarity at corresponding post-transition times (Fig. 2A, bottom). Across all ROIs, the BOLD response amplitude, averaged from 0 to 180 s after each transition, showed a strong negative correlation between LOC and ROC (r =-0.78; Fig. 2C).

To determine whether these diverse regional responses could be summarized by a small number of canonical activity patterns, we performed k-means clustering on all 77 ROIs using their concatenated BOLD time courses surrounding LOC and ROC (Fig. 2D–F). This analysis revealed four canonical spatiotemporal motifs with the highest mean silhouette score and clear spatial organization (Fig. S3). Each cluster is characterized by a distinct temporal profile (Fig. 2D) and anatomical distribution (Fig. 2E, F).

Two motifs captured the dominant polarity-reversed component of the transition response but differed markedly in their temporal kinetics. The fast association cortex–thalamic motif (C1; 29 ROIs) comprised 93% of thalamic ROIs and 71% of both polymodal association cortex and midbrain ROIs. It exhibited a rapid decrease in BOLD activity immediately after LOC and a corresponding increase after ROC. In contrast, the slow dorsal hippocampal– brainstem–cerebellar motif (C2; 16 ROIs), which included all cerebellar ROIs and 75% of both brainstem and dorsal hippocampal ROIs, displayed the opposite polarity. BOLD activity increased gradually after LOC and decreased gradually after ROC, with changes persisting throughout much of the post-transition period.

The remaining two motifs were characterized by more pronounced direction-dependent dynamics. The sensorimotor cortex–striatal motif (C3; *n* = 22), comprising 90% of sensorimotor cortex and 86% of basal ganglia ROIs, exhibited only modest changes following LOC but a prominent transient increase immediately after ROC, followed by an undershoot and gradual recovery toward baseline. In contrast, the ventral hippocampal motif (C4; *n* = 10), which contained all ventral hippocampal/parahippocampal ROIs together with smaller numbers of dorsal hippocampal, limbic, and thalamic ROIs, showed a modest transient increase after LOC but a sharp decrease immediately after ROC, followed by a pronounced rebound to, or slightly above, baseline.

Clustering performed separately on LOC-and ROC-aligned data produced highly similar patterns (Fig. S4), supporting the robustness of the identified motifs.

These analyses collectively demonstrate that regional BOLD responses during consciousness transitions are organized into a small number of reproducible spatiotemporal motifs. While most brain systems exhibit highly reversible activity patterns between LOC and ROC, a subset displays pronounced direction-dependent dynamics, indicating that consciousness transitions comprise both symmetric and asymmetric components distributed across the brain.

### Region-to-region communication exhibits reversible changes during LOC and ROC

To determine whether communication between brain regions is reversibly modulated during consciousness transitions, we quantified changes in functional connectivity (ΔFC) throughout the brain during LOC and ROC. Functional connectivity (FC) between each pair of ROIs was computed as the Pearson correlation coefficient of ROI time courses within time windows before (−360 to −60 s) and after (60 to 360 s) each transition. ROI-level ΔFC values were subsequently averaged within and between anatomical systems to generate brain system-level ΔFC matrices (Fig. 3A), in which each matrix element represents the mean connectivity change between the corresponding systems. Statistically significant ΔFC values are indicated by black dots (Fig. 3A, one-sample t-tests, FDR-corrected, q < 0.05).

**Figure 3.**
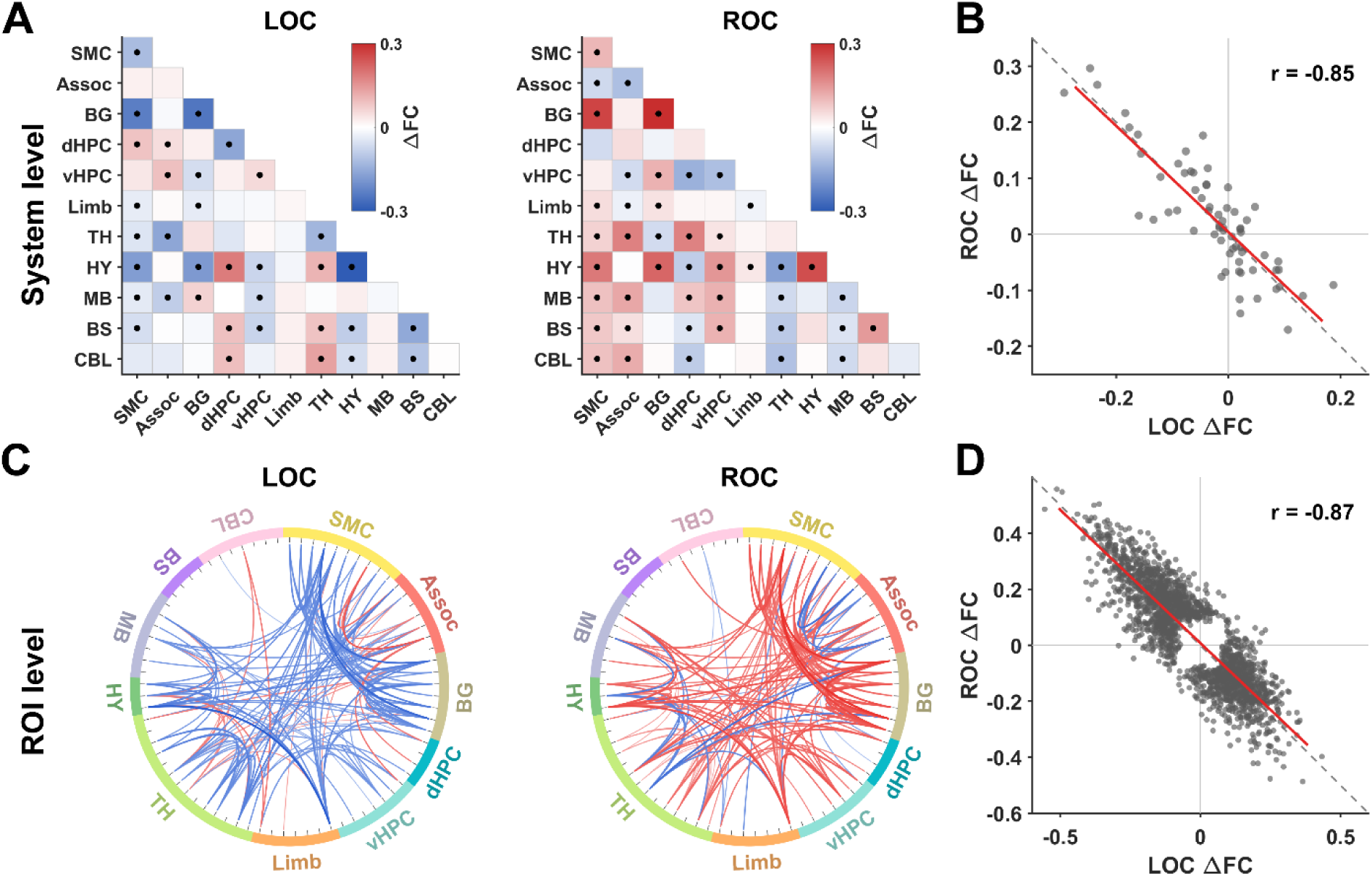
Reversed functional connectivity changes during loss and recovery of consciousness. **(A)** Functional connectivity change (ΔFC) across anatomical systems during LOC and ROC. Each cell represents the averaged ΔFC between ROIs belonging to the indicated systems. Black dots indicate system-pair ΔFC values that statistically differed from zero across scans after FDR correction (one sample t-tests, q < 0.05). Abbreviation: SMC, sensorimotor cortex; Assoc, polymodal association cortex; BG, basal ganglia; dHPC, dorsal hippocampus; vHPC, ventral hippocampal–parahippocampal system; Limb, non-hippocampal limbic system; TH, thalamus; HY, hypothalamus; MB, midbrain; BS, brainstem; CBL, cerebellum. **(B)** Relationship between system-level ΔFC during LOC and ROC. Each point represents one system–system pair. The gray dashed line denotes exact polarity reversal (i.e. ROC ΔFC = −LOC ΔFC), the red line shows the linear fit, and r is the Pearson correlation coefficient. **(C)** ROI-level connectomes showing the union of the top 100 FDR-significant ROI–ROI edges ranked by |ΔFC| separately for LOC and ROC. Red and blue curves indicate increased and decreased FC, respectively; curve width scales with |ΔFC| using common LOC/ROC scaling. Colored outer arcs denote anatomical systems. **(D)** Relationship between LOC and ROC ΔFC across the union of all ROI–ROI edges exhibiting significant ΔFC during either transition. Each point represents one ROI pair. The gray dashed line denotes exact polarity reversal, the red line shows the linear fit, and r is the Pearson correlation coefficient.

Across brain systems, connectivity changes during recovery closely mirrored those observed during LOC. System-level ΔFC patterns during LOC and ROC were strongly anticorrelated (r = −0.85; Fig. 3B), indicating that FC increases and decreases during LOC were largely reversed during ROC. This relationship remained highly robust when the analysis was applied to the collection of ROI pairs exhibiting significant ΔFC during either LOC or ROC (q < 0.05, FDR corrected; r = −0.87; Fig. 3D). Connectome visualizations of the strongest significant edge changes, using the union of the top 100 edges from LOC and ROC, further illustrate the broadly opposite pattern of FC change between the two transitions (Fig. 3C). This result remains robust when the analysis is applied to all ROI pairs (Fig. S5).

Sliding-window analysis (60-s window, 1-s step) further revealed rapid, directionally opposite FC changes around LOC and ROC across representative system pairs (Fig. S6). Together, these results demonstrate that pairwise functional connectivity is dominated by polarity-reversed changes between LOC and ROC, paralleling the largely reversible dynamics observed in local neural activity and regional BOLD responses.

### Global brain activity coordination follows distinct trajectories during LOC and ROC

Having established that local neural activity, regional BOLD responses, and inter-regional FC are largely reversible during consciousness transitions, we next asked whether the coordination of global brain activity was similarly reversible or exhibited additional path-dependent structure. We addressed this question using three complementary analyses: traveling-wave propagation, low-dimensional network trajectories and graph-theoretical organization.

### Traveling-wave propagation reveals direction-dependent organization

We first characterized traveling-wave propagation of transition-related BOLD activity across the whole brain using two independent approaches: the principal delay (PD) method based on regional peak latency^26^ and a Hilbert phase–based method^27^ referenced to the global brain signal.

The PD approach^26^ captures the timing of the dominant local BOLD deflection and extracts the most consistent spatial delay pattern across scans. Fig. 4A shows the PD profiles during LOC (left) and ROC (right). For each anatomical system, boxes represent the interquartile range of ROI-wise PD values, and systems are ordered according to their median delay. Both transitions exhibited a well-defined propagation sequence across brain systems. However, the ordering of this sequence differed substantially between LOC and ROC, resulting in distinct spatial patterns of brain activity at equivalent latencies relative to the transition points (Fig. 4A, bottom).

**Figure 4.**
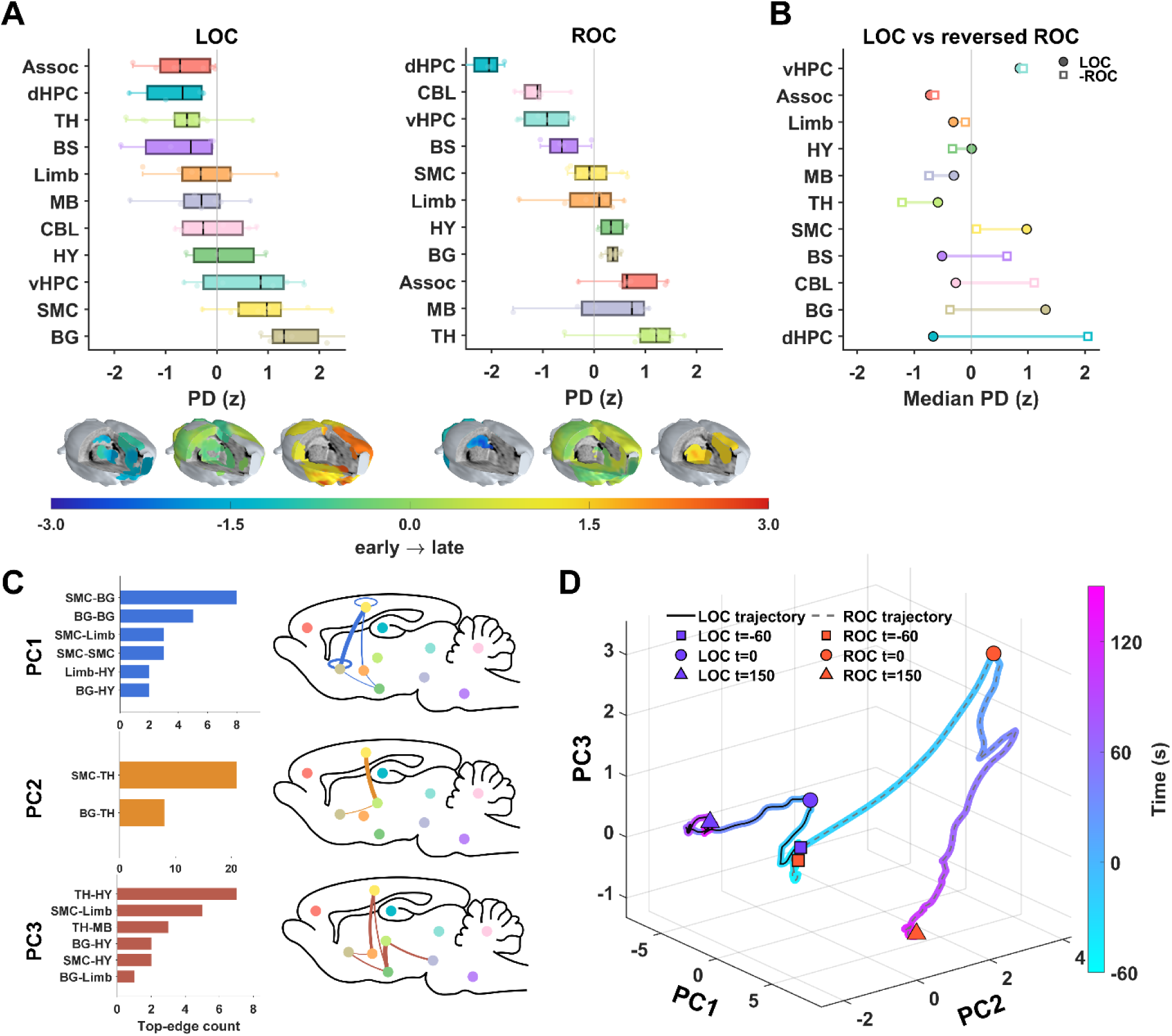
Principal-delay ordering and dominant FC trajectories during LOC and ROC. **(A)** System-level principal-delay ordering during LOC and ROC. For each system, boxes show the interquartile range of ROI-wise PD values, the vertical line inside each box indicates the median, and whiskers show the full range. ROI-level values are shown as light dots. Systems are ordered from early to late based on the median principal-delay value. Below each rank plot, representative 3D delay-bin maps show early, intermediate, and late principal-delay regions. Abbreviation: SMC, sensorimotor cortex; Assoc, polymodal association cortex; BG, basal ganglia; dHPC, dorsal hippocampus; vHPC, ventral hippocampal–parahippocampal system; Limb, non-hippocampal limbic system; TH, thalamus; HY, hypothalamus; MB, midbrain; BS, brainstem; CBL, cerebellum. **(B)** Delay-order hysteresis between LOC and ROC. The dumbbell plot compares each system’s median delay during LOC with the reversed ROC delay value. If ROC were a mirror reversal of LOC, the two markers for each system would overlap. Larger separation indicates stronger deviation from reverse symmetry. **(C)** Top enriched system pairs contributing to the first three common FC principal components. Bars show the number of top-loading FC edges assigned to each system pair for PC1, PC2, and PC3. Schematic brain maps summarize the anatomical distribution of the dominant system-pair contributions for each PC. PC1 was enriched mainly in sensorimotor–basal ganglia and other cortical–subcortical interactions, PC2 was dominated by sensorimotor–thalamic and basal ganglia–thalamic interactions, and PC3 showed additional contributions from thalamic, limbic, and cortical–subcortical system pairs. **(D)** Three-dimensional trajectory of FC dynamics during LOC and ROC projected into the shared PC1–PC2–PC3 space. Trajectories are shown from −60 to +150 s and are color-coded continuously by time relative to the pupil-defined transition. Solid black and dashed gray overlays distinguish between LOC and ROC, respectively. Squares mark −60 s, circles mark t = 0, and triangles mark +150 s relative to the corresponding transition time.

To directly evaluate the reversibility of these propagation patterns, we compared the median PD of each brain system during LOC with the sign-reversed value during ROC. If ROC simply retraced the trajectory of LOC, these two values would coincide. Instead, substantial separations were observed for multiple brain systems (Fig. 4B), demonstrating that whole-brain activity propagation deviates markedly from simple reverse symmetry.

As a complementary measure of regional temporal ordering, the Hilbert phase–based arrival-time analysis estimates when each ROI reaches the phase of the global signal and is more sensitive to the timing of the broader slow fluctuation. The phase-based analysis also revealed organized but non-reversed system-level sequences during LOC and ROC (Fig. S7A, B). Although the precise ordering of individual systems differed between the two approaches, consistent with their sensitivity to different temporal features, both analyses reached the same central conclusion: the temporal organization of brain-wide activity during ROC was not the reverse of that during LOC.

### Time-resolved functional connectivity follows distinct trajectories through network state space

To further characterize global network dynamics, we applied principal component analysis (PCA) to time-resolved FC edges during LOC and ROC. System-enrichment analysis of the top 1% contributing edges (Fig. 4C) showed that the first principal component (PC1) was primarily associated with sensorimotor–basal ganglia and other cortical–subcortical interactions, whereas PC2 was dominated by sensorimotor–thalamic and basal ganglia– thalamic connections. PC3 captured additional interactions involving thalamic, limbic, and cortical–subcortical systems.

Projection of FC dynamics into the shared PC1–PC2–PC3 space revealed both polarity-reversed and path-dependent components (Fig. 4D and Fig. S8). Along PC1, LOC and ROC moved in nearly opposite directions after the transition, and this separation persisted throughout the post-transition period. In contrast, PC2 and PC3 captured path-dependent dynamics: ROC exhibited rapid and pronounced shifts along both components, whereas LOC showed a delayed change along PC2 and only a modest change along PC3. Taken together, LOC and ROC traversed distinct trajectories through network state space as they approached their respective transition points, which can also be observed by their increasing distance over time in the space (Fig. S9).

### Global network topological organization exhibits distinct transition dynamics

We next asked whether LOC and ROC traversed distinct transition-related network configurations. Mean positive FC strength showed a sharp transient peak immediately after pupil-defined ROC, superimposed on a slower post-transition increase, whereas LOC was followed by a gradual decrease without a comparable peak (Fig. 5A). The ROC-associated FC peak coincided with an abrupt reconfiguration of global network topology. Global clustering coefficient exhibited a pronounced peak, whereas global efficiency showed a corresponding trough (Fig. 5B, C). Characteristic path length and neighborhood overlap likewise increased abruptly, while participation coefficient and local efficiency decreased (Fig. S10). In contrast, LOC showed no comparable transition-centered excursion but a sustained post-transition shift (Fig. 5B,C, Fig. S10).

**Figure 5.**
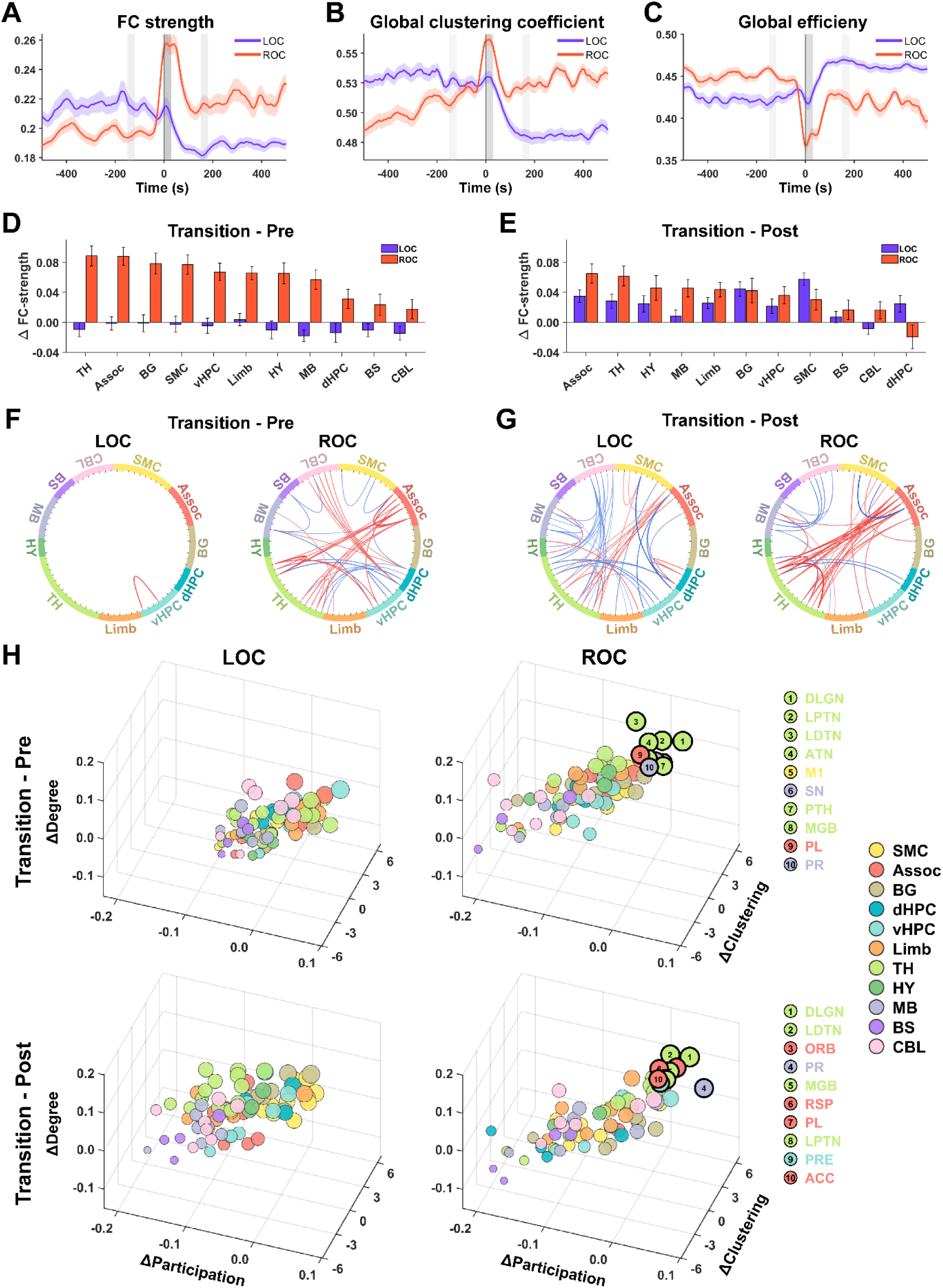
Recovery of consciousness traverses a transient high-coupling and topologically distinct network state. (**A–C**) Time courses of mean positive FC strength (A), global clustering coefficient (B), and global efficiency (C), aligned to pupil-defined LOC and ROC. FC and graph metrics were calculated using 60-s sliding windows advanced in 1-s steps; each estimate is plotted at the center of its window. Graph metrics were calculated from undirected binary networks retaining the strongest 15% of positive FC edges in each window. Lines and shaded regions show the mean and SEM across recording sessions. Gray bands mark the pre (−150 to −120 s), transition (0–30 s), and post (150–180 s) epochs of window centers. (**D,E**) Anatomical-system changes in nodal positive FC strength for transition–pre (D) and transition–post (E). Bars and error bars show the mean and SEM across recording sessions. Systems are ordered separately in each panel by descending ROC change. (**F,G**) Significant edge-occupancy changes for transition–pre (F) and transition–post (G). Edge occupancy is the fraction of windows within an epoch in which an ROI pair is retained in the fixed-density binary graph. Red and blue curves indicate increased and decreased occupancy, respectively, during the transition epoch; curve intensity and width scale with the magnitude of the occupancy change. Colored ring segments denote anatomical systems, and tick marks denote individual ROIs. **(H)** ROI-level transition-reorganization plots for transition–pre (top) and transition–post (bottom). ROIs are positioned by changes in participation coefficient, clustering coefficient, and degree; marker color denotes anatomical system, and marker size represents the sum of the three z-scored changes. Black outlines and numeric labels identify the ten highest-scoring ROC ROIs; keys at right list their abbreviations. DLGN, dorsal lateral geniculate nucleus; LPTN, lateral posterior thalamic nucleus; LDTN, laterodorsal thalamic nucleus; ATN, anterior thalamic nuclei; M1, primary motor cortex; SN, substantia nigra; PTH, posterior thalamic nucleus; MGB, medial geniculate body; PL, prelimbic cortex; PR, perirhinal cortex; ORB, orbital cortex; RSP, retrosplenial cortex; PRE, presubiculum; ACC, anterior cingulate cortex.

To characterize transition-centered network dynamics, graph metrics were estimated using a sliding-window approach (60-s window, 1-s step) and averaged within pre (−150 to −120 s), transition (0–30 s), and post (150–180 s) epochs based on window centers. During ROC, positive FC strength was higher during the transition epoch than during both comparison epochs across most anatomical systems, with the largest differences consistently observed in the thalamus and polymodal association cortex (Fig. 5D, E). Edge-occupancy maps showed the same anatomical emphasis, with prominent positive changes within and between these two systems in both ROC contrasts (Fig. 5F, G). In contrast, LOC showed little transition–pre difference but positive transition–post differences across many systems, together with more extensive transition–post edge reorganization, indicating that its network changes developed primarily after the transition.

To identify ROIs that most strongly expressed the ROC-specific topology, we characterized each ROI using transition-related topological changes in degree, participation coefficient, and nodal clustering coefficient. These three measures were z-scored across ROIs and summed to generate a descriptive transition reorganization score. The highest-scoring ROC ROIs were concentrated in thalamic and association cortex regions in both contrasts, whereas LOC showed no comparably concentrated anatomical pattern (Fig. 5H). For ROC transition–pre, six of the ten highest-ranked ROIs were thalamic nuclei (DLGN, LPTN, LDTN, ATN, PTH, and MGB), together with M1, SN, PL, and PR. For ROC transition–post, four of the ten highest-ranked ROIs were thalamic nuclei (DLGN, LDTN, MGB, and LPTN), accompanied by PR, PRE, and four association cortical regions (ORB, RSP, PL, and ACC). System-level summaries showed a similar pattern, with the thalamus and association cortex exhibiting large increases in degree and clustering together with comparatively small decreases in participation. In contrast, LOC showed relatively small transition–pre changes but larger and more broadly distributed transition–post changes in these nodal metrics (Fig. S11).

Taken together, these findings show that ROC traverses a transient, distributed high-coupling state characterized by greater local clustering and reduced global integration, with its strongest expression in thalamic and association regions. LOC lacked a comparable transition-centered state and instead underwent a slower, more broadly distributed post-transition reorganization.

These three complementary analyses all demonstrate that the global evolution of network organizations followed distinct paths during LOC and ROC.

### Distinct global trajectories are not explained by neuromodulatory drive

Because pupil dynamics provide a sensitive measure of arousal-related brainstem neuromodulatory tone, we next asked whether the observed differences in global brain dynamics could simply reflect differences in autonomic arousal. For each ROI, we quantified the cross-correlation between pupil diameter and BOLD activity over the interval from −60 to 180 s surrounding each transition.

In contrast to the pronounced differences observed in traveling-wave propagation, network trajectories, and brain topology, brain-wide BOLD–pupil coupling was remarkably similar during LOC and ROC. The spatial distribution of coupling strength was highly conserved between the two transitions (Fig. 6A), with strong correlations in both coupling magnitude (r = 0.82; Fig. 6B) and temporal lag (Fig. 6C) across ROIs. These findings indicate that neuromodulatory influences on brain-wide BOLD activity are largely preserved across LOC and ROC. Therefore, the distinct trajectories of global brain organization cannot be simply attributed to differences in neuromodulatory tone, but instead likely arise from direction-dependent reconfiguration of large-scale brain networks.

**Figure 6.**
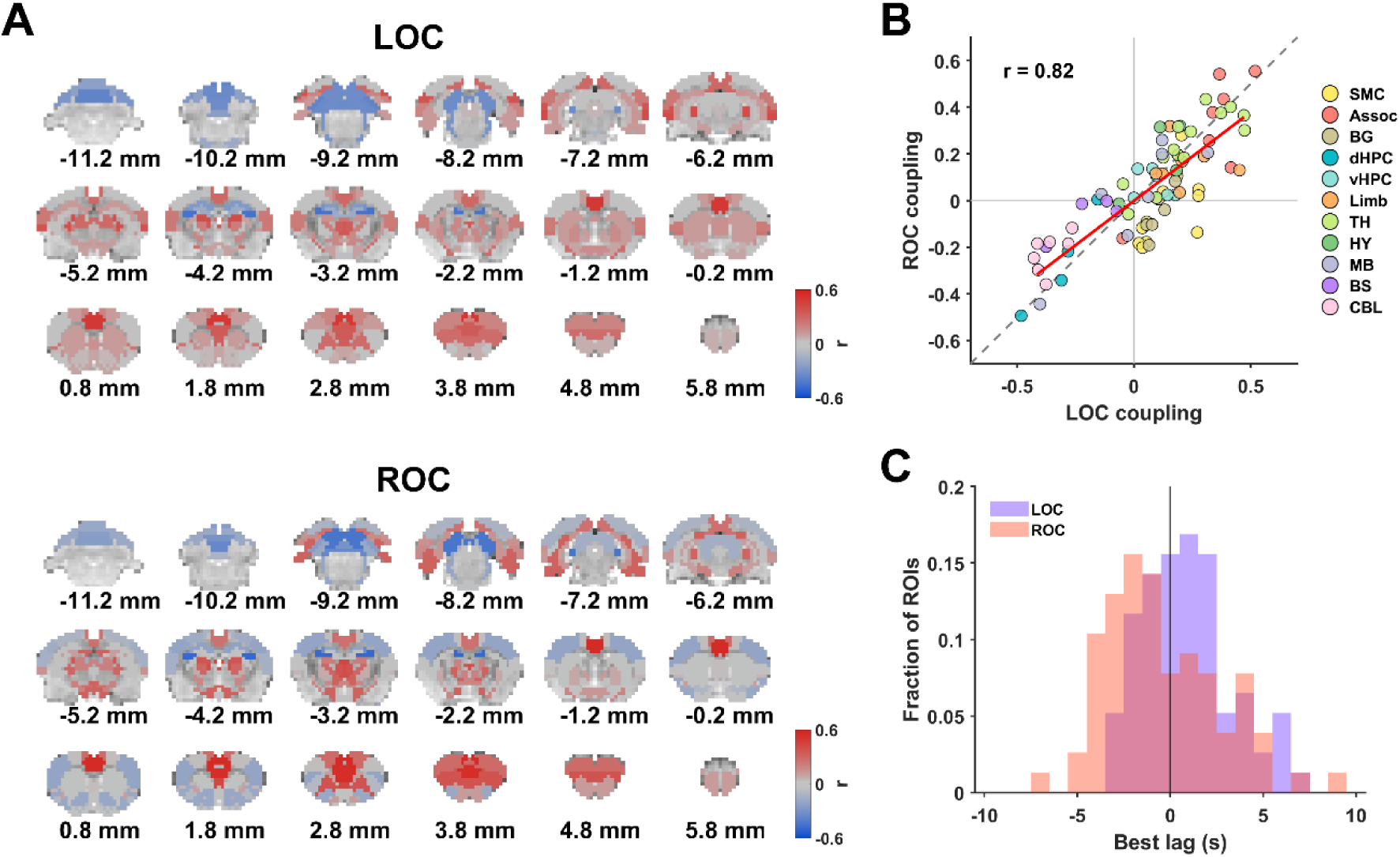
Pupil–BOLD coupling dynamics during loss and recovery of consciousness. **(A)** Spatial distribution of pupil–BOLD timecourse coupling during LOC and ROC. For each ROI, pupil and BOLD signals were correlated over the −60 to 180 s window relative to LOC or ROC across lags from −10 to +10 s. The correlation at the lag with the largest absolute value was retained with its original sign and mapped onto coronal slices. **(B)** Relationship between LOC and ROC pupil–BOLD coupling across ROIs. Each dot represents one ROI, colored by anatomical system. The positive correlation indicates that regions with stronger pupil–BOLD coupling during LOC tend to show similar coupling strength during ROC. Abbreviation: SMC, sensorimotor cortex; Assoc, polymodal association cortex; BG, basal ganglia; dHPC, dorsal hippocampus; vHPC, ventral hippocampal–parahippocampal system; Limb, non-hippocampal limbic system; TH, thalamus; HY, hypothalamus; MB, midbrain; BS, brainstem; CBL, cerebellum. **(C)** Distribution of ROI-wise best lags for pupil–BOLD coupling during LOC and ROC. Histograms show the lag at which the absolute pupil–BOLD correlation was maximal for each ROI, summarizing the temporal offset between pupil dynamics and regional BOLD responses. Purple: LOC; orange: ROC; red: overlap.

Collectively, multiple independent analyses—including traveling-wave propagation, low-dimensional network trajectories, and graph-theoretical organization—consistently revealed direction-dependent reorganization of global brain dynamics during LOC and ROC. Importantly, these differences cannot be explained by changes in neuromodulatory tone, as brain-wide pupil–BOLD coupling remains highly consistent across the two transitions. These results demonstrate that brain activity during consciousness transitions is locally reversible but globally path-dependent, reflecting intrinsic reconfiguration of large-scale brain networks rather than differences in autonomic arousal.

## Discussion

By integrating local electrophysiology, whole-brain imaging, and autonomic measures within a unified experimental framework, this study provides a multiscale characterization of neural dynamics around consciousness transitions modulated by anesthesia. Our results reveal a consistent and principled dissociation across scales: while local brain activities, including electrophysiology signals, regional BOLD activities as well as FC of individual connections are largely reversible between LOC and ROC, the global coordination of brain activity follows distinct trajectories during the two transitions. These results are illustrated in Fig. 7. Our data suggest that although the level of pharmacological suppression can decide the local brain response, the transition into and out of unconsciousness is governed by history-dependent reconfiguration of large-scale brain networks.

**Figure 7.**
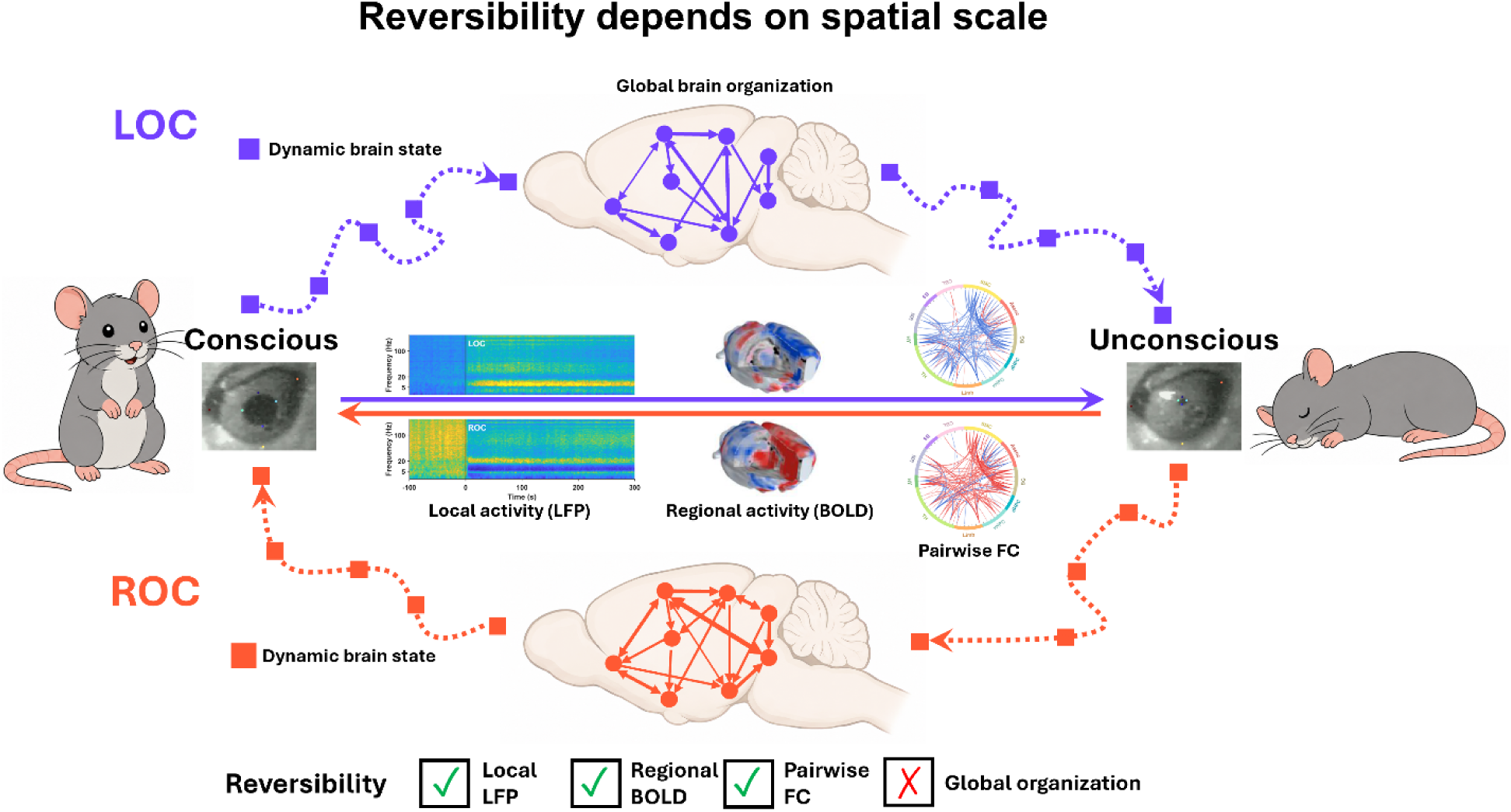
Summary schematic of brain dynamics during loss and recovery of consciousness. LOC (purple) and ROC (orange) showed largely reversible changes in pupil dynamics, regional brain activity, and functional connectivity, whereas their global network trajectories did not retrace each other, indicating direction-dependent reorganization of whole-brain activity.

At the local level, we observed robust increases in LFP power during LOC and corresponding decreases during ROC in both the secondary motor cortex and dorsal hippocampus, consistent with previous studies demonstrating structured and reversible oscillatory changes during propofol anesthesia^4,5,16^. These electrophysiological changes were tightly synchronized with pupil constriction and dilation, respectively, indicating close coupling between local neural activity and autonomic arousal. Because pupil diameter covaries with locus coeruleus activity and reflects cholinergic, parasympathetic, and other neuromodulatory processes^22–24^, these observations establish pupil dynamics as a temporally precise physiological marker of consciousness transitions. In addition, regional BOLD activity and pairwise FC exhibited highly symmetric changes between LOC and ROC. Together, these findings indicate that neural responses at local and inter-regional levels are largely reversible and closely reflect the graded pharmacological effects of anesthesia.

Despite this overall reversibility, temporal patterns of regional BOLD responses were not spatially homogeneous. K-means clustering revealed four reproducible spatiotemporal motifs, suggesting that brain regions sharing similar transition dynamics form functionally coordinated networks during consciousness transitions. The association cortex–thalamus motif is particularly notable. The thalamus contains both core and matrix projection systems, with matrix neurons projecting broadly to association cortex and coordinating distributed cortical activity. Recent work indicates that consciousness-related thalamic effects are nucleus-specific, with distinct roles for core-and matrix-type systems across pharmacological and pathological unconsciousness^28,29^. Human fMRI studies have shown that propofol preferentially disrupts interactions between matrix-rich thalamic regions and higher-order association cortex^13^, consistent with our observation that these regions form a coherent functional motif during LOC and ROC. Previous neuroimaging studies likewise demonstrated preferential disruption of higher-order thalamocortical and frontoparietal networks, whereas primary sensory systems remained relatively preserved^30,31^.

The sensorimotor–basal ganglia motif exhibited a distinct response profile. These structures are linked through parallel cortico-striato-pallido-thalamocortical loops that regulate movement initiation, action selection, and behavioral gating^32^. Their relatively modest responses during LOC are consistent with preservation of low-level sensory and motor processing after higher-order cortical integration has already deteriorated. In contrast, the dorsal hippocampus–brainstem–cerebellum motif exhibited BOLD changes opposite to those of the association cortex–thalamus motif, whereas the ventral hippocampus displayed a unique transient suppression during ROC before recovering toward baseline. These distinct temporal signatures are consistent with the well-established functional and anatomical differences between dorsal and ventral hippocampus. Collectively, these observations indicate that consciousness transitions are organized at the level of functional brain systems rather than isolated anatomical regions.

While regional activity is largely reversible, our analyses reveal a fundamentally different picture at the global level. Traveling-wave analysis demonstrated that brain-wide activity propagates across distributed systems in an organized manner during both LOC and ROC, but with markedly different propagation sequences. In particular, both the principal-delay and phase-based analyses showed that the association cortex–thalamus system occupies an early position during LOC but a late position during ROC. This finding has a plausible biological interpretation. Experimental studies have shown that higher-order thalamic activity changes before widespread cortical activity during transitions into sleep and propofol anesthesia^33,34^. Our results therefore support a model in which disruption of higher-order thalamocortical recurrence represents one of the earliest systems-level events during LOC. Conversely, the late recruitment of this network during ROC suggests that restoration of higher-order thalamocortical interactions is among the final steps required for stable recovery of conscious processing. This interpretation is consistent with studies showing early subcortical and sensorimotor recruitment during awakening, followed by delayed reactivation of higher-order cortical networks^35,36^.

The remaining brain systems do not simply reverse their temporal ordering. Sensorimotor cortex and basal ganglia occupy relatively late positions during LOC but become engaged much earlier during ROC, consistent with early restoration of motor readiness and behavioral responsiveness. Similarly, transient involvement of brainstem and hypothalamic systems during recovery agrees with recruitment of ascending arousal pathways reported in human propofol studies^37^. The dorsal hippocampus and cerebellum likewise exhibit propagation patterns that deviate from reverse symmetry, likely reflecting region-specific synchronization and autonomic regulation rather than a single common mechanism.

These observations suggest a two-stage model of consciousness transitions. During LOC, higher-order association cortex and thalamic systems reorganize first, followed by progressive engagement of hippocampal, cerebellar, subcortical, and finally sensorimotor– basal ganglia systems as the brain settles into the anesthetized state. During ROC, this sequence is not simply reversed. Instead, arousal-related subcortical and sensorimotor systems become re-engaged early, whereas restoration of higher-order thalamocortical organization occurs later, ultimately re-establishing the large-scale coordination required for conscious processing.

The traveling-wave results are reinforced by two independent analyses of global network organization. Trajectories of functional connectivity projected into a shared low-dimensional state space showed that LOC and ROC traverse distinct paths as they approach their respective transition points.

Graph metric analysis further showed that recovery of consciousness is not simply the temporal reversal of loss of consciousness. During its progression toward a more segregated and functionally differentiated brain network, ROC was marked by a sharp, coordinated transient excursion across multiple network metrics, followed by a return toward the slower evolving trend. Specifically, positive FC strength, global clustering, and characteristic path length showed transient peaks, whereas global efficiency and mean participation coefficient showed corresponding troughs. The abrupt increase in functional coupling was not accompanied by greater global integration; instead, the strongest connections were transiently redistributed into more locally closed and overlapping neighborhoods and concentrated within a narrower set of anatomical-system relationships. These changes suggest that recovery transiently overshoots toward a more differentiated network configuration, with coupling becoming unusually local and system-specific, before the balance between segregation and integration characteristic of the recovered state was re-established.

Although this reorganization was distributed across multiple anatomical systems, it was most prominent in the thalamus and polymodal association cortex. These systems showed the largest and most consistent changes in FC strength, edge occupancy, and nodal topology across both transition–pre and transition–post contrasts. In particular, greater edge occupancy within and between thalamic and association regions indicates that these connections formed a prominent component of the transient strongest-edge configuration. The convergent system-, edge-, and ROI-level findings therefore identify a preferentially coupled thalamic–association coalition as the most prominent anatomical feature of the ROC transient state. In contrast, LOC lacked a comparable intermediate network state, and instead showed a gradual post-transition shift toward weaker coupling and greater global integration.

Importantly, the divergent global trajectories during transitions cannot be explained simply by changes in neuromodulatory drive. Our pupil–BOLD analysis suggests that neuromodulatory influences remain remarkably similar during LOC and ROC. Therefore, the distinct global trajectories are unlikely to arise from different neuromodulatory drives themselves, but rather from the fact that comparable neuromodulatory inputs act upon fundamentally different large-scale network reconfigurations.

Taken together, our findings suggest that the hysteresis observed during LOC and ROC emerges primarily at the level of large-scale network organization rather than local neural activity. They support the view that recovery of consciousness is not simply the reversal of anesthetic induction, but an active process of large-scale network reorganization. These results are also aligned with computational frameworks in which conscious states are metastable dynamical configurations whose transition paths need not be symmetric^38^.

Several limitations should be acknowledged. Although propofol provides a highly controlled model for studying reversible unconsciousness, it remains unclear whether the principles identified here generalize to other anesthetic agents or to physiological and pathological alterations of consciousness, including natural sleep and disorders of consciousness. Future studies combining broader electrophysiological sampling, causal perturbation techniques, and whole-brain imaging across multiple states will be important for determining the generality of the multiscale organization identified here.

In summary, our findings demonstrate that consciousness transitions are characterized by a coexistence of reversible local neural dynamics and path-dependent global network organization. This multiscale dissociation provides new insight into the systems-level mechanisms of anesthesia-induced unconsciousness and suggests that recovery of consciousness represents an active reassembly of large-scale brain networks rather than simply the reversal of pharmacological suppression. These findings have important implications for understanding consciousness, improving anesthetic monitoring, and interpreting neural state transitions across both physiological and pathological conditions.

## Methods

### Animals

Twelve adult male Long–Evans rats (400–550 g) were used in this study. Animals were housed in Plexiglas cages under controlled temperature (22–24 °C) on a 12-h light/dark cycle with food and water available *ad libitum*. All experimental procedures were approved by the Pennsylvania State University Institutional Animal Care and Use Committee (IACUC protocol #PRAMS201343583) and were conducted in accordance with the National Institutes of Health Guide for the Care and Use of Laboratory Animals.

### Surgical procedures

For electrophysiological recordings, animals underwent stereotaxic implantation of MR-compatible electrodes. Anesthesia was induced with ketamine (40 mg kg⁻¹) and xylazine (12 mg kg⁻¹) and maintained with 0.5–1.0% isoflurane delivered through a nose cone throughout the procedure. Body temperature was maintained using a feedback-controlled heating pad (PhysioSuite, Kent Scientific), while heart rate and arterial oxygen saturation were continuously monitored using a pulse oximeter (MouseSTAT^®^ Jr, Kent Scientific).

A linear 16-channel MR-compatible silicon probe (A1×16-5mm-150-177-MRCM16LP, NeuroNexus) was implanted unilaterally to target the dorsal hippocampus and secondary motor cortex (M2). Stereotaxic coordinates relative to bregma were AP −3.7 mm, ML −1.5 mm, DV 3.0 mm for the hippocampus, and AP +2.6 mm, ML −1.5 mm, DV 1.2 mm for M2. Reference and ground electrodes consisted of silver wires placed on the contralateral cerebellar surface. The implant was secured with dental cement (Metabond, Parkell).

Following surgery, animals received subcutaneous enrofloxacin (Baytril, 2.5 mg kg⁻¹) and long-acting buprenorphine (1.0 mg kg⁻¹) and were allowed to recover for at least one week before imaging experiments.

### Simultaneous fMRI, pupillometry, respiration and electrophysiology recordings

Rats were positioned in a custom-built restraint apparatus compatible with simultaneous MRI, electrophysiology, pupillometry, and respiration recordings. An MR-compatible infrared camera was positioned lateral to the right eye for continuous pupil monitoring. To maintain eyelid opening, a lightweight plastic ring was attached to the right eyelid using silicone adhesive (KWIK-SIL, World Precision Instruments), and ophthalmic lubricant was applied to prevent corneal dehydration. Core body temperature was maintained at 36.5– 37.5 °C using a circulating warm-water system.

Propofol was administered through a tail-vein catheter using a stepwise infusion protocol. A 20 mg kg⁻¹ propofol bolus was first administered through a tail-vein catheter. After allowing at least 30 min for the animal to reach a stable anesthetic state, continuous multimodal recording was initiated, and propofol was infused in a stepwise manner at 20 mg kg⁻¹ h⁻¹ for 20 min, followed by 40 mg kg⁻¹ h⁻¹ for 15 min and 80 mg kg⁻¹ h⁻¹ for 10 min. Propofol infusion was then discontinued, and recordings continued for an additional 35 min during spontaneous recovery, yielding a total recording duration of 80 min (Fig. 1C).

MRI experiments were performed on a 7-T Bruker BioSpec 70/30 scanner (ParaVision 6.0.1, Bruker) equipped with a laboratory-built single-loop surface coil. Resting-state fMRI data were acquired using a T2*-weighted gradient-echo echo-planar imaging (EPI) sequence (TR = 1000 ms, TE = 15 ms, slice thickness = 1 mm, in-plane resolution = 0.5 × 0.5 mm², 20 slices, field of view = 32 × 32 mm², matrix size = 64 × 64, 4800 volumes per scan). Structural images were acquired using a T2-weighted rapid acquisition with relaxation enhancement (RARE) sequence (TR = 3000 ms, TE = 40 ms, matrix size = 256 × 256, field of view = 32 × 32 mm², slice thickness = 1 mm, 20 slices, six averages).

Electrophysiological signals were acquired simultaneously using a Tucker-Davis Technologies (TDT) recording system. Local field potentials were recorded through an MR-compatible 16-channel headstage (LP16CH), amplified and digitized with a PZ5 neurodigitizer, processed by an RZ2 BioAmp processor, and sampled at 24,414 Hz. Data were recorded using Synapse software (TDT).

Pupil diameter was recorded at 30 frames s⁻¹ using an MR-compatible infrared camera (MR-CAM, model 12M_i, MRC Systems GmbH) and synchronized to MRI acquisition via transistor–transistor logic (TTL) pulses generated by an Arduino microcontroller running custom Python software. Animals breathed spontaneously throughout the experiment. Respiration was monitored using a thoracic pressure sensor positioned beneath the chest (sampling rate = 225 Hz) and synchronized to MRI acquisition through scanner-generated TTL pulses.

### fMRI data preprocessing

fMRI data were preprocessed using our previously published pipeline^39^, including removal of the first 10 volumes, motion scrubbing based on framewise displacement (FD > 0.15 mm), spatial smoothing (Gaussian kernel, FWHM = 1 mm), nuisance regression of motion parameters and white matter and cerebrospinal fluid (CSF) signals, independent component analysis (ICA)-based artifact removal, and temporal band-pass filtering (0.01– 0.1 Hz).

### Electrophysiological data preprocessing

Multichannel electrophysiology data acquired simultaneously with fMRI were preprocessed to remove MRI-induced artifacts using a dynamic template regression pipeline^4,17,20^. Raw electrophysiological signals were first temporally aligned to the fMRI acquisition by identifying recurrent transients associated with slice acquisition. Based on this alignment, MRI-induced interference was estimated and removed separately for each slice window. Slice onset times were initially detected from transient interference peaks and subsequently refined by aligning slice-locked electrophysiological segments to an averaged interference template using cross-correlation. Slice-specific interference templates were then constructed by averaging aligned segments within a sliding window spanning 400 neighboring slice acquisitions, thereby capturing slow temporal variations in interference amplitude and waveform. For each slice window, the corresponding template was temporally aligned to the raw signal using cross-correlation and removed by linear regression. The regression residual was retained as the cleaned electrophysiological signal. Following template regression, power-line contamination was attenuated using notch filters centered at 60 Hz and its harmonics. Residual periodic interference at the slice acquisition frequency (20 Hz) and its harmonics was further removed using narrow band-stop filters. The cleaned signal was subsequently band-pass filtered (0.1–300 Hz, zero-phase Butterworth filter) to obtain the local field potential (LFP). To further suppress residual high-frequency structured noise, slice-locked signal segments were arranged into matrices and subjected to principal component analysis (PCA). The leading principal components were then regressed from the signal within each slice window, thereby reducing residual variance not removed by the template regression procedure.

Time-frequency power was estimated from the denoised LFP using a multitaper spectrogram implemented in MATLAB (*mtspecgramc*)^40^. Multitaper spectrograms were computed using a 3-s window with a 0.1-s step, a time–bandwidth product of 3, five tapers, and a frequency range of 0.35–300 Hz. To minimize contamination from motion-related artifacts, time points corresponding to excessive head motion, defined by FD, were excluded from the analysis. Spectrogram power at each frequency bin was converted to percent change relative to the pre-transition baseline (ΔP/P, %) using the −180 to −30 s interval relative to LOC or ROC. Band-limited power time courses were obtained by averaging ΔP/P values across frequency bins within eight canonical frequency bands: slow wave (SW, 0.35–1 Hz), delta (1–4 Hz), theta (4–8 Hz), alpha (8–12 Hz), beta (12–30 Hz), low gamma (30–50 Hz), middle gamma (50–100 Hz), and high gamma (100–180 Hz).

For group-level analyses, outliers across scans were removed at each time-frequency point using a median absolute deviation (MAD)-based robust z-score (|Z| ≥ 3). Band-limited ΔP/P time courses were then smoothed using a 5-s moving-average filter. To quantify post-transition spectral changes, the largest absolute deviation from baseline within the 0–360 s post-transition window was identified for each frequency band, and the signed ΔP/P value at that time point was retained.

### Respiration data preprocessing

Respiration peaks and troughs were identified from the thoracic pressure trace using a threshold-based peak detection algorithm. Respiration rate (breaths min⁻¹) was calculated from successive peak-to-peak intervals.

### Pupil data preprocessing

Pupil videos were cropped to a fixed region of interest and contrast-enhanced to improve visualization of the pupil boundary. Pupil landmarks were then tracked using DeepLabCut^21^, and pupil diameter and center position were computed from the tracked coordinates using an in-house MATLAB pipeline. Frames with low tracking confidence or blinks were excluded, and brief tracking failures were corrected by interpolation. For a small subset of frames, pupil landmarks were manually labeled to improve tracking accuracy. The resulting pupil diameter time series was synchronized to the fMRI acquisition using recorded TTL triggers. For group-level analyses, each pupil time course was normalized by its within-session standard deviation before transition alignment and averaging.

### Determination of LOC and ROC

Loss and recovery of consciousness were determined from pupil diameter dynamics. State transitions were identified based on sustained changes in pupil diameter and its first derivative. LOC was defined as the onset of rapid, sustained pupil constriction preceding the anesthetized plateau, characterized by a prominent negative peak in the first derivative within a predefined temporal window. ROC was defined analogously as the onset of rapid, sustained pupil dilation following the anesthetized plateau, characterized by a prominent positive peak in the first derivative. For both transitions, candidate events were required to satisfy predefined derivative amplitude and absolute pupil diameter thresholds to exclude transient fluctuations. All automatically detected transition points were visually inspected and verified.

To assess cross-modal temporal agreement, LFP-defined LOC and ROC were independently identified as the time points corresponding to the steepest increase and decrease, respectively, in alpha-band power. Signed temporal offsets were calculated as the LFP-defined transition time minus the pupil-defined transition time. Pupil-defined LOC and ROC were used as the common temporal reference for all subsequent multimodal analyses.

### BOLD signal normalization and voxel-wise dynamics around transitions

BOLD time courses were converted to percent signal change (ΔBOLD%) relative to the pre-transition baseline (−180 to −30 s relative to LOC or ROC. Whole-brain BOLD signals were summarized using a custom 77-ROI parcellation derived from the Waxholm Space rat brain atlas^41^, with ROIs grouped into 11 anatomical systems. Unless otherwise specified, all subsequent analyses were performed on ΔBOLD% time courses aligned to the transition point. To visualize whole-brain activity dynamics during LOC and ROC, voxel-wise ΔBOLD% volumes spanning −100 to +300 s relative to the transition point were extracted and compiled into Supplementary Movies 1 and 2.

### ROI time course clustering

For each ROI, voxel-wise ΔBOLD% signals were averaged to obtain ROI time courses, which were subsequently averaged across scans. Scan-averaged ROI responses within the −30 to +150 s peri-transition window were clustered using correlation-distance *k*-means clustering. Candidate solutions with *k* = 2–8 were evaluated using 50 random initializations and a maximum of 2,000 iterations. The four-cluster solution, which yielded the highest mean silhouette score, was selected for all subsequent analyses. Cluster-average time courses were computed by averaging the ROI time courses within each cluster. For visualization, each ROI’s cluster label was assigned to its atlas-defined spatial mask.

### Functional connectivity analysis

For each scan, FC between every pair of ROIs was quantified as the Pearson correlation between ROI BOLD time courses within the pre-transition (−360 to −60 s) and post-transition (60 to 360 s) windows. ΔFC were defined as post-transition FC minus pre-transition FC. ROI-pair ΔFC values were then averaged within and between anatomical systems to generate system-level connectivity matrices. Statistical significance of ΔFC relative to zero was assessed across scans and false discovery rate (FDR) corrected. System-level connectivity matrices were visualized as lower triangular matrices, including the within-system diagonal elements. Reversibility between LOC and ROC was quantified as the Pearson correlation between the corresponding entries of the two system-level ΔFC matrices.

Time-resolved FC was estimated using a sliding-window approach with a 60-s window advanced in 1-s increments. For selected system pairs, sliding-window FC was calculated by averaging pairwise correlations between ROIs belonging to the corresponding anatomical systems. The resulting FC time courses were then averaged across scans and are reported as mean ± SEM.

### Traveling-wave propagation analysis

Traveling-wave propagation timing across ROIs was estimated using two complementary approaches applied to transition-aligned ROI ΔBOLD% time courses across scans. Missing time points resulting from motion scrubbing were linearly interpolated to obtain continuous peri-transition time courses.

### Principal delay method

Using a peak-latency–based approach^26^, a delay profile was computed for each scan by identifying the peak time of each ROI within the peri-transition window (−15 to +15 s) and expressing each peak time as a relative delay with respect to the transition time. After concatenating delay profiles across scans, the resulting ROI-by-scan delay matrix was decomposed using singular value decomposition (SVD). The first left singular vector defined the principal delay (PD) profile, representing the dominant spatial ordering of response delays across ROIs. PD values were mapped onto the brain atlas for visualization.

For system-level analyses, ROI PD values were grouped according to anatomical system. Systems were ordered by their median PD, and the median and interquartile range were displayed together with the underlying ROI-level values. Reverse symmetry was evaluated by comparing each system’s median PD during LOC with the sign-reversed median PD during ROC. A value of (−ROC) − LOC equal to zero indicated perfect reversibility.

### Instantaneous phase method

The global signal (GS) was computed as the mean ΔBOLD% across all voxels within the brain mask. ROI and GS time courses were mildly smoothed using a Gaussian kernel (width = 3). Instantaneous phase was estimated using the Hilbert transform followed by phase unwrapping^27^. The direction of GS phase evolution was determined from the sign of the mean temporal derivative of the unwrapped GS phase within a ±10 s window centered on the transition.

For each ROI, the arrival time was defined as the phase zero-crossing closest to the transition within a ±10 s search window, constrained to have the same crossing direction as the GS. Arrival times were expressed relative to the transition time. Group-level arrival times were obtained by averaging ROI arrival-time vectors across scans. Arrival times were mapped onto the brain atlas for visualization, and reverse symmetry was evaluated analogously by comparing system median arrival times during LOC with the sign-reversed medians during ROC.

### Principal component analysis of time-resolved functional connectivity

For each scan, sliding-window FC matrices were computed as described above. FC values were Fisher *z*-transformed prior to analysis. Each FC matrix was vectorized by extracting the upper triangular elements, yielding a time-by-edge matrix representing the temporal evolution of FC. Analyses were restricted to the peri-transition window (−360 to +360 s relative to the transition). To reduce inter-scan variability, FC edges were baseline-corrected on a per-scan basis by subtracting, for each edge, its mean FC during the pre-transition baseline period (−360 to −60 s) from all peri-transition windows within the same scan. Missing values resulting from motion scrubbing were replaced using edge-wise mean values computed across all available windows in the pooled dataset.

To identify low-dimensional trajectories of FC dynamics during LOC and ROC, PCA was performed on time-resolved FC edges pooled across both transitions. A pooled training matrix was constructed by concatenating all valid peri-transition windows from LOC and ROC scans, with rows corresponding to time windows and columns corresponding to FC edges. The pooled matrix was column-centered and decomposed using SVD. The resulting right singular vectors defined orthogonal principal components in edge space, representing the dominant patterns of FC covariance shared between LOC and ROC. Deriving the PCA basis from the pooled dataset ensured that both transitions were represented within a common low-dimensional subspace, thereby enabling direct comparison of their trajectories.

To facilitate interpretation, FC edges were ranked according to the absolute loading magnitude within each retained principal component, and the top 1% of edges were identified. System-level enrichment was quantified by counting the number of top-loading edges associated with each anatomical system, thereby summarizing the systems contributing most strongly to each principal component.

Finally, baseline-corrected FC edge vectors from each scan were projected onto the common PCA basis to obtain principal component score time courses. Mean trajectories and standard errors were then computed across scans for each principal component. To visualize the dominant low-dimensional FC dynamics, LOC and ROC trajectories were plotted in the three-dimensional PC1–PC2–PC3 space over the interval from −60 to +150 s, with time color-coded along each trajectory to illustrate the temporal evolution of FC states across the transition.

### Graph theoretical analysis

Time-resolved FC matrices were obtained from the 77-ROI BOLD time series using a 60 s sliding window advanced in 1 s steps and aligned to pupil-defined LOC or ROC. FC strength was calculated directly from these weighted FC matrices as the mean positive connectivity across all unique ROI pairs. Graph-theoretical analyses were performed using the Brain Connectivity Toolbox in MATLAB^42^. For graph construction, the diagonal elements of each FC matrix were removed, and the top 15% of positive connections were retained to generate undirected binary graphs (network density = 0.15).

### Clustering coefficient

For node *i*, degree was calculated as the number of edges incident on the node. The nodal clustering coefficient was then calculated as:

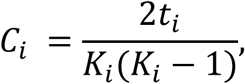

where *t_i_* is the number of triangles containing node *i*, and *K_i_* is the degree of node *i*. Nodes with fewer than two neighbors were assigned a clustering coefficient of zero. The global clustering coefficient was defined as the mean nodal clustering coefficient across all ROIs.

### Characteristic path length and global efficiency

Characteristic path length was calculated as the mean shortest-path distance across all node pairs. Global efficiency was calculated as the average inverse shortest-path length across all node pairs.

### Local efficiency

For each node, local efficiency was calculated as the global efficiency of the subgraph formed by its immediate neighbors after excluding the node itself. Mean local efficiency was obtained by averaging nodal values across all ROIs.

### Neighborhood overlap

For each connected ROI pair, neighborhood overlap was calculated as the normalized number of neighbors shared by the two endpoints. Mean neighborhood overlap was obtained by averaging this value across all graph edges.

### Participation coefficient

The participation coefficient of node *i* was defined as

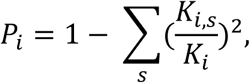

where *K_i_*_,*s*_ is the number of edges linking node *i* to its anatomical system *s*, and *K_i_* is the total degree of node *i*. Here, s indexes the 11 predefined anatomical systems. Mean participation coefficient was calculated by averaging nodal participation coefficients across all ROIs.

### Edge occupancy

An edge-occupancy analysis was performed to characterize transition-centered reorganization of the binary functional network. For each scan, edge occupancy was defined as the fraction of sliding windows within each epoch (transition: 0–30 s; pre: −150 to −120 s; post: 150–180 s) in which an edge was retained in the fixed density binary graph. Occupancy differences were calculated separately as transition-pre and transition-post. For each edge, occupancy differences were tested across scans using one-sample t-tests against zero, followed FDR correction across all ROI pairs.

### Transition reorganization score

Degree, participation coefficient and clustering coefficient were calculated for each ROI in every sliding window. To identify ROIs showing coordinated changes across multiple nodal graph metrics, a transition reorganization score was calculated separately for the transition–pre and transition–post contrasts:

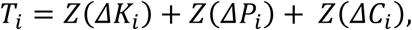

where Δ denotes the difference between the transition epoch and the corresponding pre-or post-transition epoch. Thus, *T_i_* represents the sum of the standardized changes in degree (*K_i_*), participation coefficient (*P_i_*) and clustering coefficient for ROI *i*.

### Pupil–BOLD coupling analysis

To test whether direction-dependent global trajectories could be explained by differences in arousal-related drive, pupil diameter was cross-correlated with the BOLD time course of each ROI over the interval from −60 to +180 s relative to pupil-defined LOC and ROC. Cross-correlations were evaluated over lags from −10 to +10 s. For each ROI and transition, the peak cross-correlation coefficient and its corresponding temporal lag were extracted. The spatial distributions of coupling strength and temporal lag were then compared between LOC and ROC across ROIs using Pearson correlation.

## Acknowledgments

The authors express their gratitude to Dr. Thomas Neuberger for his assistance and guidance in the MRI facility, to Dr. Patrick Drew for his constructive scientific discussion, and to Drs. Dennis C.Y. Chan and Hayreddin S. Ünsal for their technical support. This work is partially supported by the National Institutes of Health (R01GM141792 to N.Z.).

## Author contributions

N.Z. developed the concept. N.Z. and X.H. designed experiments. X.H., X.C. and S.C. set up experiments. X.H. conducted the experiments. X.H. conducted data analysis. Y.D. assisted with data analysis. X.H. prepared the figures. N.Z. secured funding. X.H. and N.Z. wrote the manuscript. X.H. and N.Z. edited the manuscript.

## References

1. Laureys, S., Owen, A. M. & Schiff, N. D. Brain function in coma, vegetative state, and related disorders. The Lancet Neurology 3, 537–546 (2004).

2. Tononi, G., Boly, M., Massimini, M. & Koch, C. Integrated information theory: from consciousness to its physical substrate. Nature Reviews Neuroscience 17, 450–461 (2016).

3. Mashour, G. A. & Hudetz, A. G. Neural Correlates of Unconsciousness in Large-Scale Brain Networks. Trends in Neurosciences 41, 150–160 (2018).

4. Chen, X., Cramer, S. R., Chan, D. C. Y., Han, X. & Zhang, N. Sequential Deactivation Across the Hippocampus-Thalamus-mPFC Pathway During Loss of Consciousness. Advanced Science 11, e2406320 (2024).

5. Purdon, P. L. et al. Electroencephalogram signatures of loss and recovery of consciousness from propofol. Proc. Natl. Acad. Sci. U.S.A. 110, (2013).

6. Lewis, L. D. et al. Rapid fragmentation of neuronal networks at the onset of propofol-induced unconsciousness. Proc. Natl. Acad. Sci. U.S.A. 109, (2012).

7. Bastos, A. M. et al. Neural effects of propofol-induced unconsciousness and its reversal using thalamic stimulation. eLife 10, e60824 (2021).

8. Xiong, Y. (Sophy) et al. Propofol-mediated loss of consciousness disrupts predictive routing and local field phase modulation of neural activity. Proc. Natl. Acad. Sci. U.S.A. 121, e2315160121 (2024).

9. Hudetz, A. G. & Mashour, G. A. Disconnecting Consciousness: Is There a Common Anesthetic End Point? Anesthesia & Analgesia 123, 1228–1240 (2016).

10. Fiset, P. et al. Brain Mechanisms of Propofol-Induced Loss of Consciousness in Humans: a Positron Emission Tomographic Study. J. Neurosci. 19, 5506–5513 (1999).

11. Stamatakis, E. A., Adapa, R. M., Absalom, A. R. & Menon, D. K. Changes in Resting Neural Connectivity during Propofol Sedation. PLoS ONE 5, e14224 (2010).

12. Akeju, O. et al. Disruption of thalamic functional connectivity is a neural correlate of dexmedetomidine-induced unconsciousness. eLife 3, e04499 (2014).

13. Huang, Z., Mashour, G. A. & Hudetz, A. G. Propofol disrupts the functional core-matrix architecture of the thalamus in humans. Nat Commun 15, 7496 (2024).

14. Sanders, R. D., Tononi, G., Laureys, S. & Sleigh, J. W. Unresponsiveness ≠ unconsciousness. Anesthesiology 116, 946–959 (2012).

15. Friedman, E. B. et al. A Conserved Behavioral State Barrier Impedes Transitions between Anesthetic-Induced Unconsciousness and Wakefulness: Evidence for Neural Inertia. PLoS ONE 5, e11903 (2010).

16. Brown, E. N., Purdon, P. L. & Van Dort, C. J. General anesthesia and altered states of arousal: a systems neuroscience analysis. Annual Review of Neuroscience 34, 601–628 (2011).

17. Tu, W. & Zhang, N. Neural underpinning of a respiration-associated resting-state fMRI network. eLife 11, e81555 (2022).

18. Tu, W., Cramer, S. & Zhang, N. Disparity in temporal and spatial relationships between resting-state electrophysiological and fMRI signals. Preprint at 10.21203/rs.3.rs-3251741/v5 (2024).

19. Turner, K. L., Gheres, K. W. & Drew, P. J. Relating Pupil Diameter and Blinking to Cortical Activity and Hemodynamics across Arousal States. J. Neurosci. 43, 949–964 (2023).

20. Tu, W., Cramer, S. R. & Zhang, N. Disparity in temporal and spatial relationships between resting-state electrophysiological and fMRI signals. eLife 13, RP95680 (2024).

21. Mathis, A. et al. DeepLabCut: markerless pose estimation of user-defined body parts with deep learning. Nat Neurosci 21, 1281–1289 (2018).

22. Murphy, P. R., O’Connell, R. G., O’Sullivan, M., Robertson, I. H. & Balsters, J. H. Pupil diameter covaries with BOLD activity in human locus coeruleus. Human Brain Mapping 35, 4140–4154 (2014).

23. Reimer, J. et al. Pupil fluctuations track rapid changes in adrenergic and cholinergic activity in cortex. Nat Commun 7, 13289 (2016).

24. Joshi, S., Li, Y., Kalwani, R. M. & Gold, J. I. Relationships between Pupil Diameter and Neuronal Activity in the Locus Coeruleus, Colliculi, and Cingulate Cortex. Neuron 89, 221–234 (2016).

25. Mukamel, E. A. et al. A transition in brain state during propofol-induced unconsciousness. The Journal of Neuroscience 34, 839–845 (2014).

26. Gu, Y. et al. Brain Activity Fluctuations Propagate as Waves Traversing the Cortical Hierarchy. Cerebral Cortex 31, 3986–4005 (2021).

27. Raut, R. V. et al. Global waves synchronize the brain’s functional systems with fluctuating arousal. Sci. Adv. 7, eabf2709 (2021).

28. Whyte, C. J., Redinbaugh, M. J., Shine, J. M. C Saalmann, Y. B. Thalamic contributions to the state and contents of consciousness. Neuron 112, 1611–1625 (2024).

29. Szocs, D. et al. A thalamic perspective of (un)consciousness in pharmacological and pathological states in humans. Brain Communications 8, fcag021 (2026).

30. Boveroux, P. et al. Breakdown of within-and between-network resting state functional magnetic resonance imaging connectivity during propofol-induced loss of consciousness. Anesthesiology 113, 1038–1053 (2010).

31. Liu, X. et al. Propofol attenuates low-frequency fluctuations of resting-state fMRI BOLD signal in the anterior frontal cortex upon loss of consciousness. NeuroImage 147, 295– 301 (2017).

32. Alexander, G. E., DeLong, M. R. & Strick, P. L. Parallel organization of functionally segregated circuits linking basal ganglia and cortex. Annual Review of Neuroscience 9, 357–381 (1986).

33. Magnin, M. et al. Thalamic deactivation at sleep onset precedes that of the cerebral cortex in humans. Proceedings of the National Academy of Sciences of the United States of America 107, 3829–3833 (2010).

34. Baker, R. et al. Altered Activity in the Central Medial Thalamus Precedes Changes in the Neocortex during Transitions into Both Sleep and Propofol Anesthesia. The Journal of Neuroscience 34, 13326–13335 (2014).

35. Alcaide, S. et al. fMRI lag structure during waking up from early sleep stages. Cortex 142, 94–103 (2021).

36. Setzer, B. et al. A temporal sequence of thalamic activity unfolds at transitions in behavioral arousal state. Nature Communications 13, 5442 (2022).

37. Nir, T. et al. Transient subcortical functional connectivity upon emergence from propofol sedation in human male volunteers: evidence for active emergence. British Journal of Anaesthesia 123, 298–308 (2019).

38. Kringelbach, M. L. & Deco, G. Brain States and Transitions: Insights from Computational Neuroscience. Cell Reports 32, 108128 (2020).

39. Liu, Y. et al. An open database of resting-state fMRI in awake rats. NeuroImage 220, 117094 (2020).

40. Bokil, H., Andrews, P., Kulkarni, J. E., Mehta, S. & Mitra, P. P. Chronux: A platform for analyzing neural signals. Journal of Neuroscience Methods 192, 146–151 (2010).

41. Kleven, H. et al. Waxholm Space atlas of the rat brain: a 3D atlas supporting data analysis and integration. Nat Methods 20, 1822–1829 (2023).

42. Rubinov, M. & Sporns, O. Complex network measures of brain connectivity: Uses and interpretations. NeuroImage 52, 1059–1069 (2010).

