## Supplementary Document for "Brain dynamics exhibit scale-dependent reversibility during consciousness transitions"

**This PDF file includes:**

Supplementary Figures 1 to 11

Captions for Supplementary Movies 1 to 2

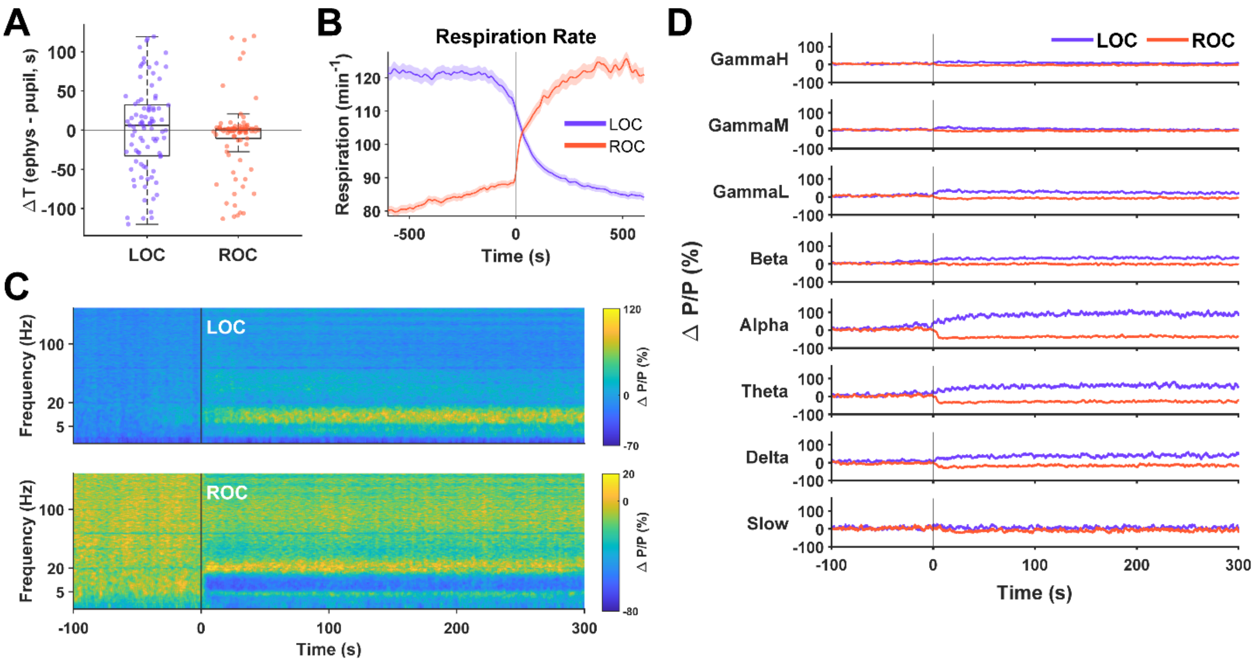


**Supplementary Figure 1. Dorsal hippocampal electrophysiology and respiration dynamics during propofol-induced LOC and ROC.**

(**A**) Timing difference between independently estimated hippocampal alpha-band LFP transitions and pupil-defined transitions. Boxplots summarize the group distributions, and each dot represents one recording session. Positive values indicate that the LFP transition occurred after the pupil transition; negative values indicate that it occurred before. The mean signed timing difference (LFP-defined minus pupil-defined) was 2.0 s for LOC (95% CI, -9.9 to 14.0 s) and -7.9 s for ROC (95% CI, -17.3 to 1.5 s).

(**B**) Respiration rate aligned to pupil-defined LOC and ROC. Lines and shaded regions show the mean and SEM across recording sessions; the vertical line marks the transition time.

(**C**) Averaged dorsal hippocampal spectrograms aligned to pupil-defined LOC and ROC. Spectral power is expressed as percent change relative to the pre-transition normalization baseline. LOC was accompanied by broadband power increases, whereas ROC showed broadly opposite changes.

(**D**) Band-limited dorsal hippocampal LFP power changes aligned to pupil-defined LOC and ROC for slow-wave, delta, theta, alpha, beta, GammaL, GammaM, and GammaH bands. Lines and shading show the mean and SEM across recording sessions. Peak post-transition ΔP/P values for LOC were 19.2%, 50.1%, 74.8%, 106.0%, 39.1%, 36.8%, 20.5%, and 15.7%, respectively; corresponding values for ROC were −21.4%, −29.7%, −37.0%, −46.1%, −7.0%, −11.8%, −5.1%, and −6.7%.

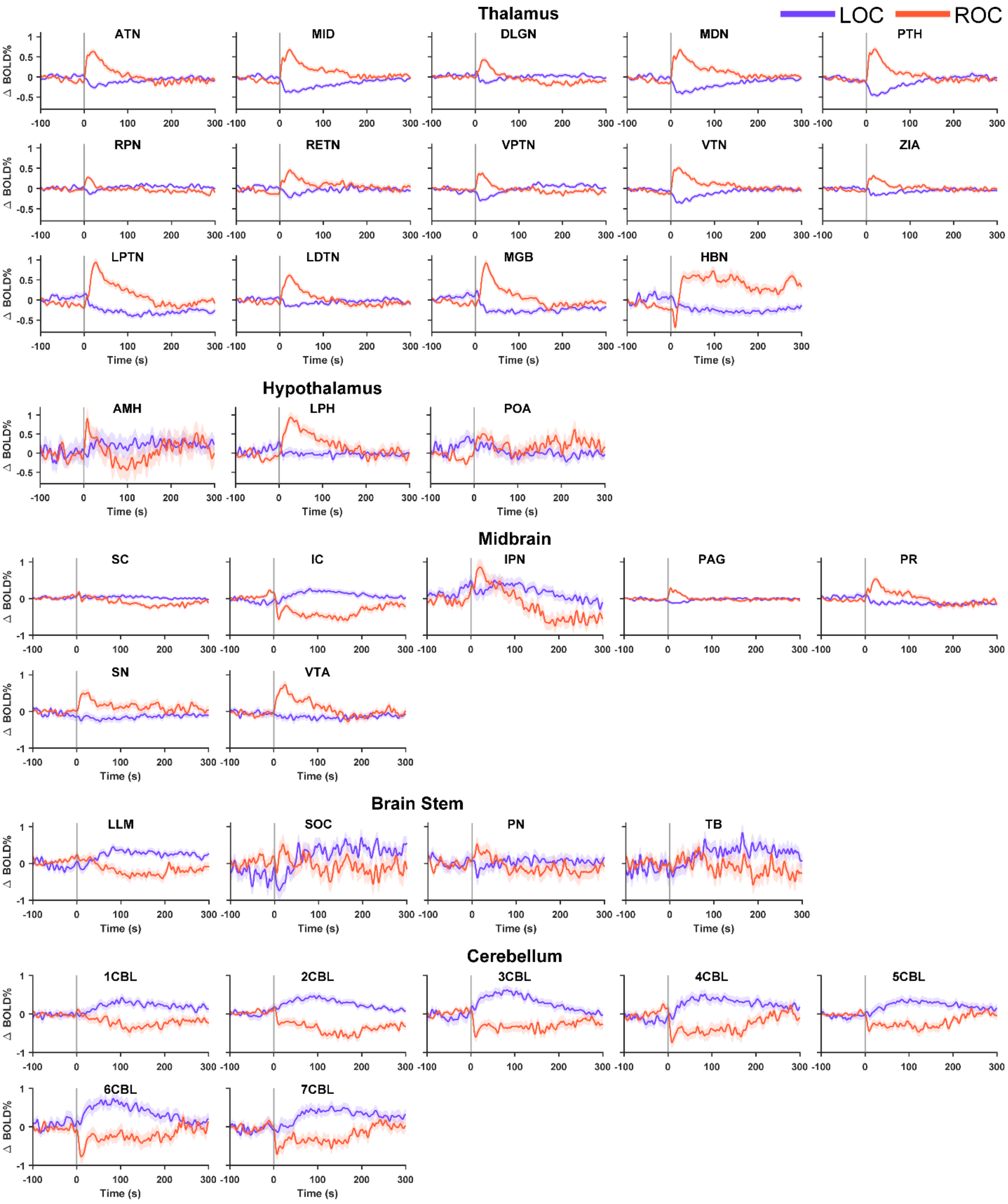

Thalamus
LOC
ROC
ATN
MID
DLGN
MDN
PTH
RPN
RETN
VPTN
VTN
ZIA
LPTN
LDTN
MGB
HBN
Hypothalamus
AMH
LPH
POA
Midbrain
SC
IC
IPN
PAG
PR
SN
VTA
Brain Stem
LLM
SOC
PN
TB
Cerebellum
1CBL
2CBL
3CBL
4CBL
5CBL
6CBL
7CBL
Δ BOLD%
Time (s)

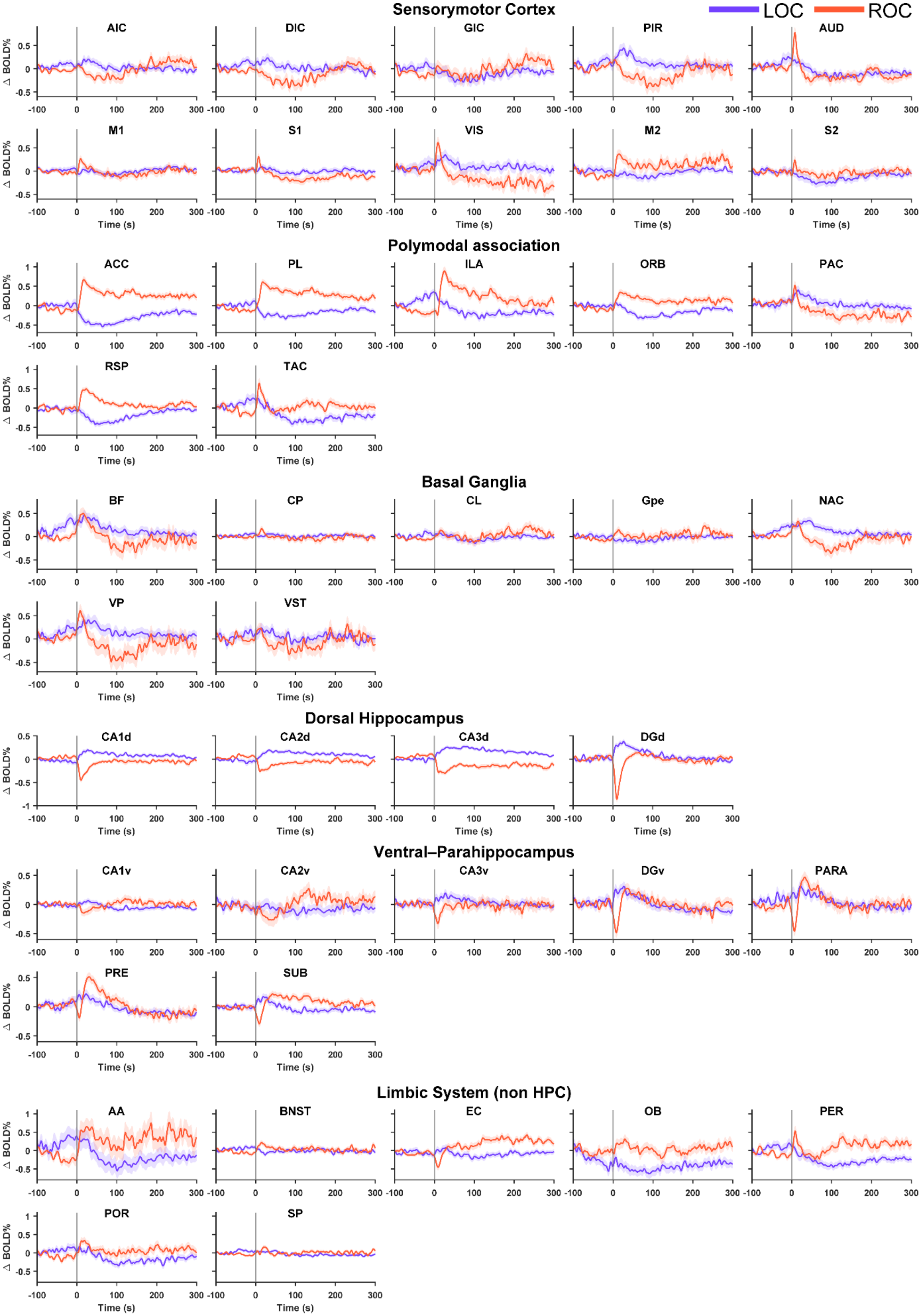

LOC
ROC
Sensorymotor Cortex
AIC
DIC
GIC
PIR
AUD
M1
S1
VIS
M2
S2
Polymodal association
ACC
PL
ILA
ORB
PAC
RSP
TAC
Basal Ganglia
BF
CP
CL
Gpe
NAC
VP
VST
Dorsal Hippocampus
CA1d
CA2d
CA3d
DGd
Ventral–Parahippocampus
CA1v
CA2v
CA3v
DGv
PARA
PRE
SUB
Limbic System (non HPC)
AA
BNST
EC
OB
PER
POR
SP
Δ BOLD%
Time (s)

**Supplementary Figure 2. ROI-level BOLD time courses during LOC and ROC.**

Mean BOLD signal changes are shown for individual ROIs grouped by the corresponding anatomical system. Purple and orange traces indicate pupil-defined LOC and ROC, respectively, aligned to the transition at 0 s. Shaded regions indicate SEM across recording sessions.

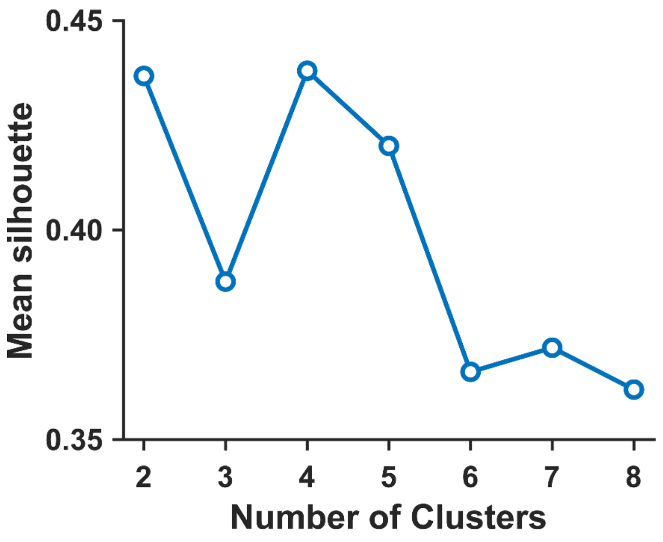


**Supplementary Figure 3. Mean silhouette scores for different cluster numbers.**

Mean silhouette scores are shown for correlation-distance k-means solutions with k = 2–8 using concatenated LOC- and ROC-aligned ROI BOLD response profiles.

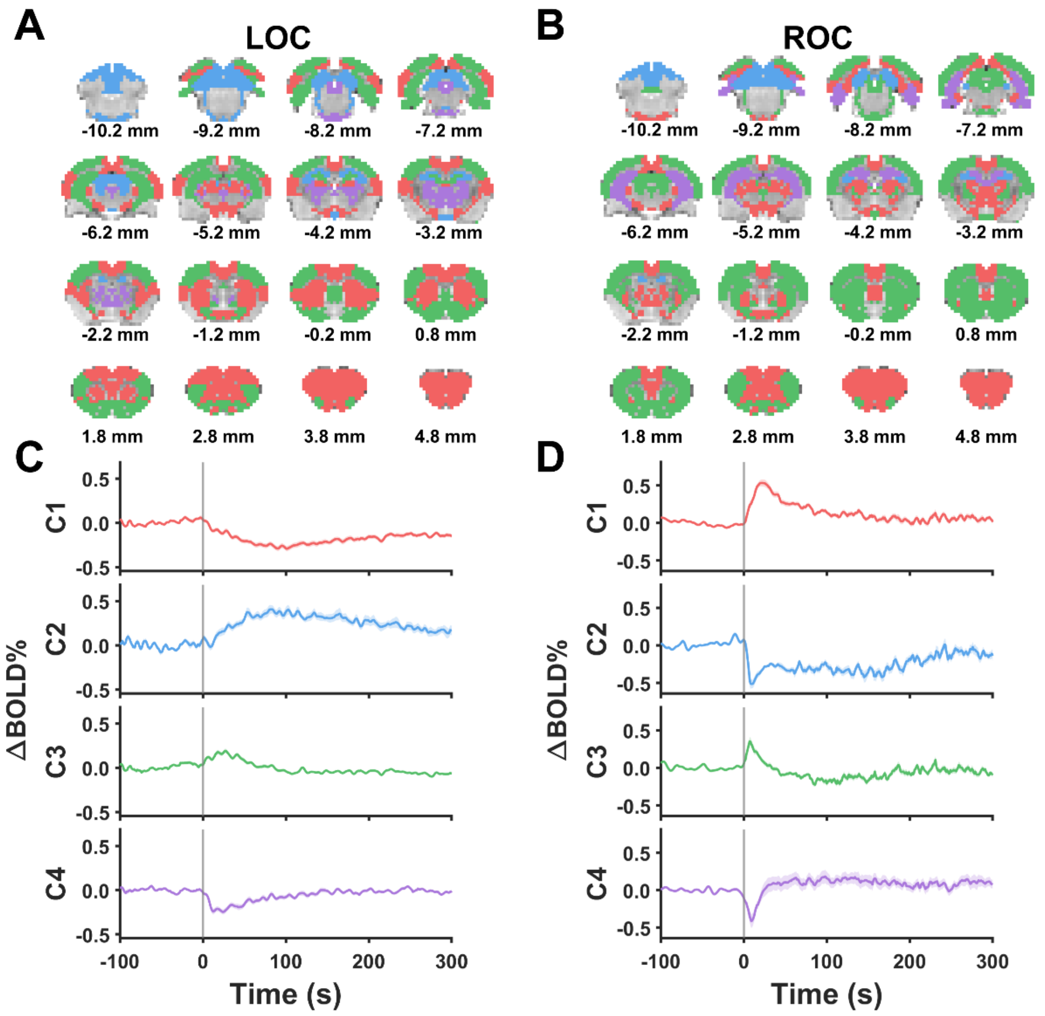


**Supplementary Figure 4. Separate clustering of ROI BOLD time courses around LOC and ROC.**

K-means clustering was performed separately on LOC- and ROC-aligned ROI BOLD time courses. (**A,B**) Coronal maps show the spatial distribution of the color-coded LOC and ROC clusters. (**C,D**) Mean BOLD time courses for each cluster, averaged across ROIs within the cluster; shading indicates SEM across ROIs.

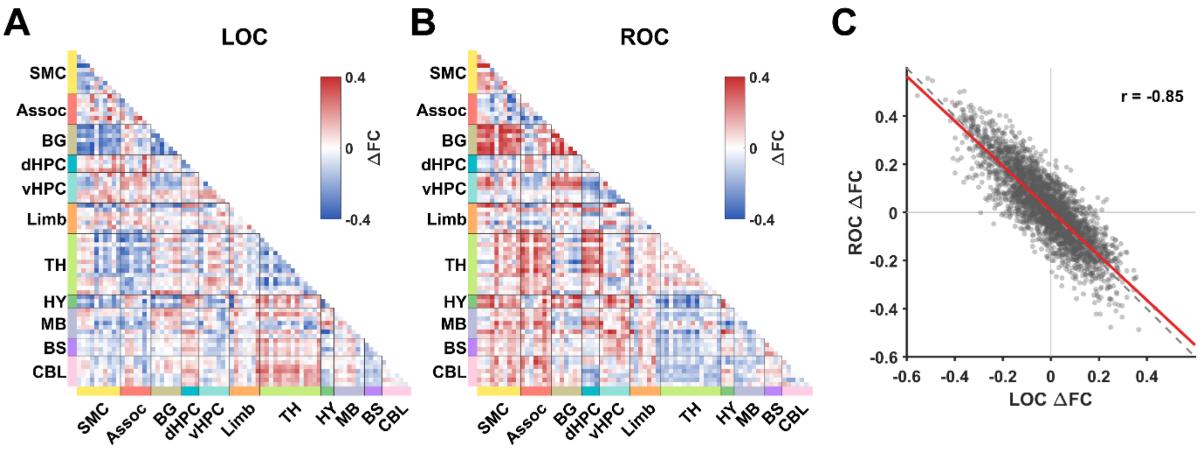


**Supplementary Figure 5. ROI-wise functional connectivity changes during LOC and ROC.**

(**A,B**) ROI-wise ΔFC matrices for LOC and ROC. ΔFC was calculated as post-transition FC (60 to 360 s) minus pre-transition FC (−360 to −60 s) using ROI-averaged BOLD signals. ROIs are grouped by anatomical system. Red and blue indicate increased and decreased FC, respectively. System abbreviations: SMC, sensorimotor cortex; Assoc, polymodal association cortex; BG, basal ganglia; dHPC, dorsal hippocampus; vHPC, ventral hippocampal/parahippocampal system; Limb, nonhippocampal limbic system; TH, thalamus; HY, hypothalamus; MB, midbrain; BS, brainstem; CBL, cerebellum.

(**C**) Relationship between LOC and ROC ΔFC across all ROI pairs. Each point represents one ROI pair. The gray dashed line denotes exact polarity reversal, the red line shows the linear fit, and r is the Pearson correlation coefficient.

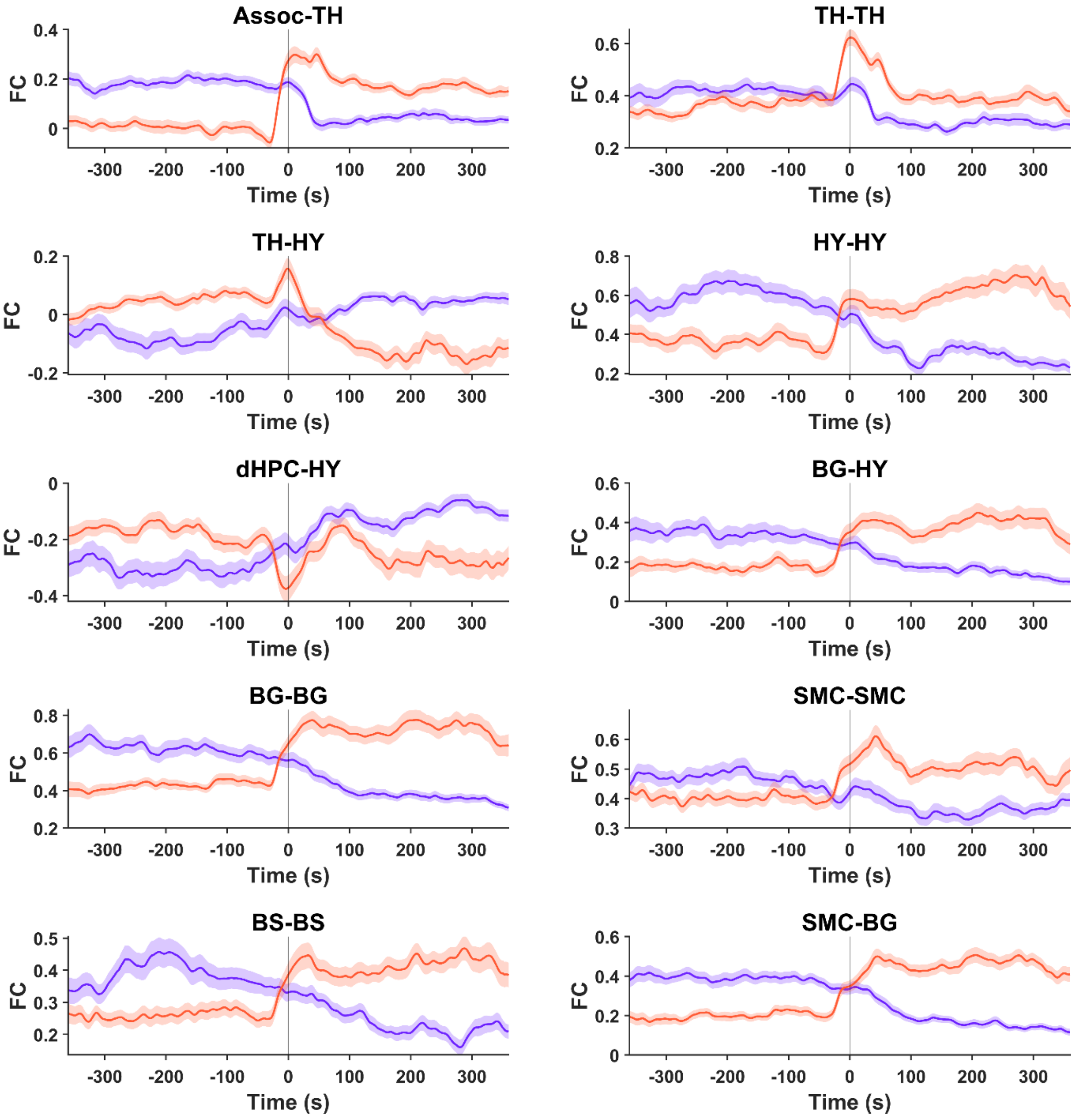


**Supplementary Figure 6. Representative system–system dynamic FC trajectories around LOC and ROC.**

Purple and orange lines indicate LOC- and ROC-aligned FC, respectively. FC was calculated using 60-s sliding windows and plotted at each window center. Lines and shaded regions show the mean and SEM across recording sessions. Time zero marks the pupil-defined transition. System abbreviations: SMC, sensorimotor cortex; Assoc, polymodal association cortex; BG, basal ganglia; dHPC, dorsal hippocampus; TH, thalamus; HY, hypothalamus; BS, brainstem.

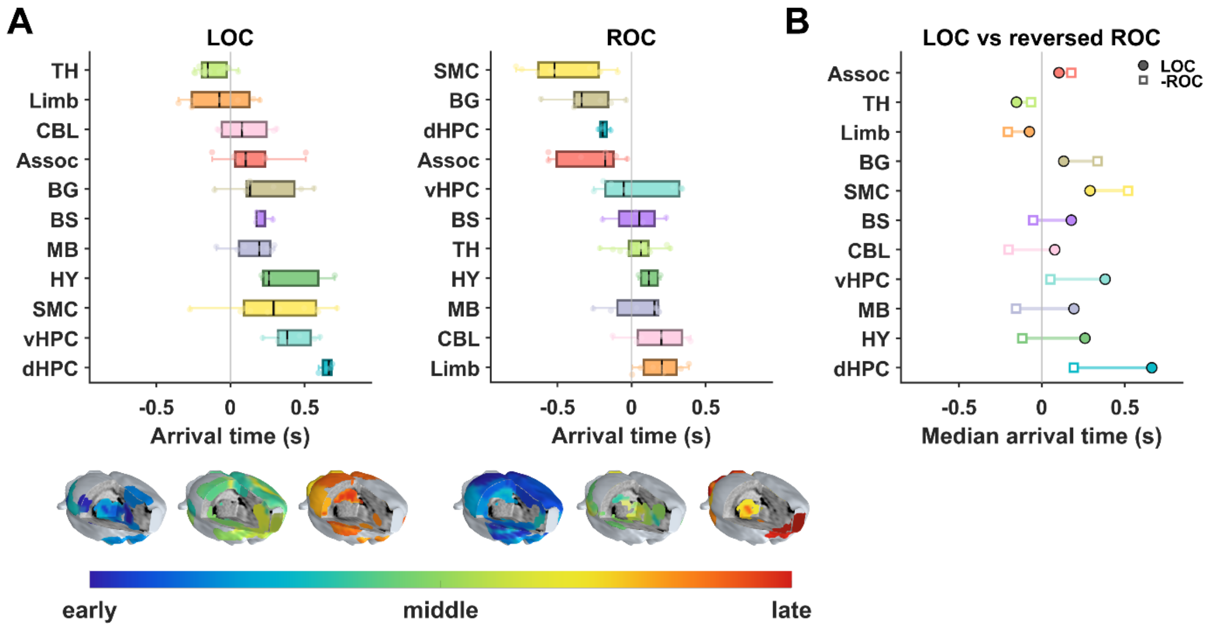


**Supplementary Figure 7. Phase-based transition-wave ordering during LOC and ROC.**

(A) System-level phase-arrival ordering during LOC and ROC. For each system, boxes show the interquartile range of ROI-wise arrival times, the internal line indicates the median, whiskers show the full range, and light dots show individual ROIs. Systems are ordered from early to late by median arrival time. Maps below show early, intermediate, and late rank-normalized delay bins; boxplots retain arrival times in seconds.

(B) Comparison of LOC phase-arrival time with sign-reversed ROC phase-arrival time. Exact overlap would indicate reverse symmetry.

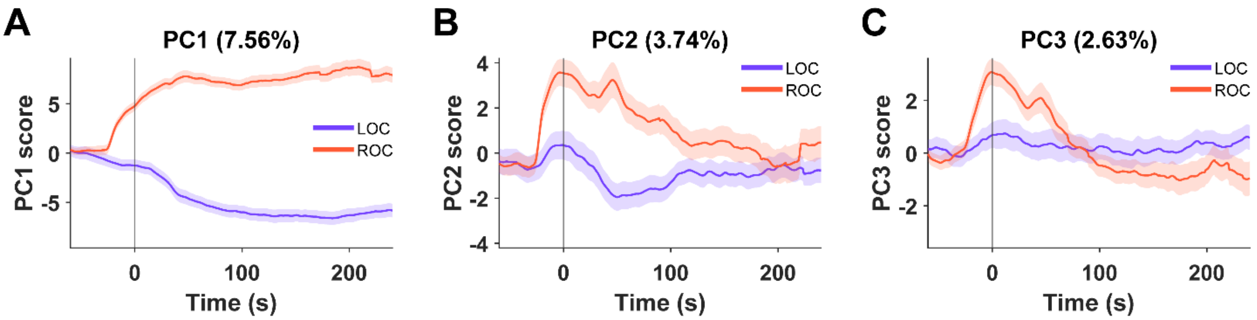


**Supplementary Figure 8. Time courses of the leading three shared FC principal components during LOC and ROC.**

(**A–C**) Lines and shaded regions show the mean PC score and SEM across recording sessions. Purple and orange indicate LOC and ROC, respectively; time zero marks the pupil-defined transition. PC1 explained 7.56% of variance and captured a sustained polarity-reversed axis. PC2 explained 3.74% and showed a transient ROC-dominant excursion followed by gradual relaxation. PC3 explained 2.63% and captured an additional early transition-asymmetric component.

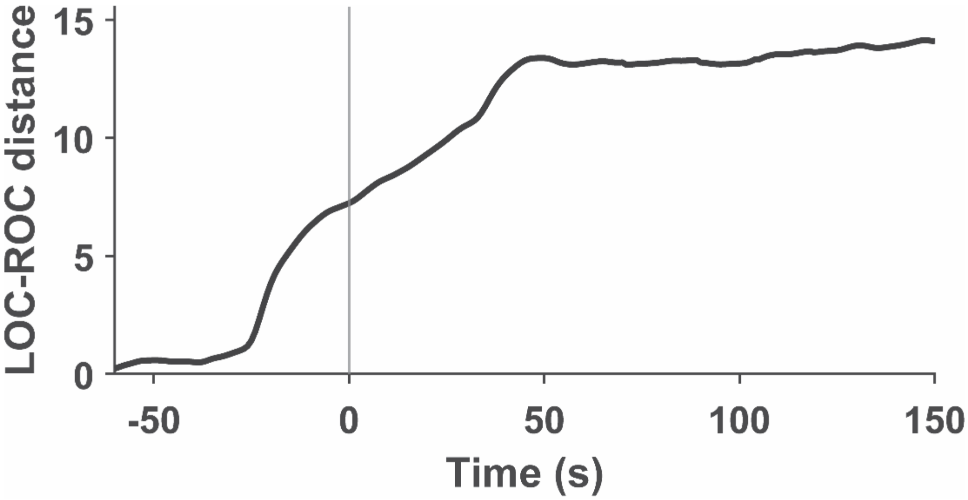


**Supplementary Figure 9. Temporal separation between LOC and ROC trajectories in the leading three-PC space.**

At each time point, Euclidean distance was calculated between the mean LOC and ROC coordinates in the shared PC1–PC2–PC3 space. The vertical line marks the pupil-defined transition.

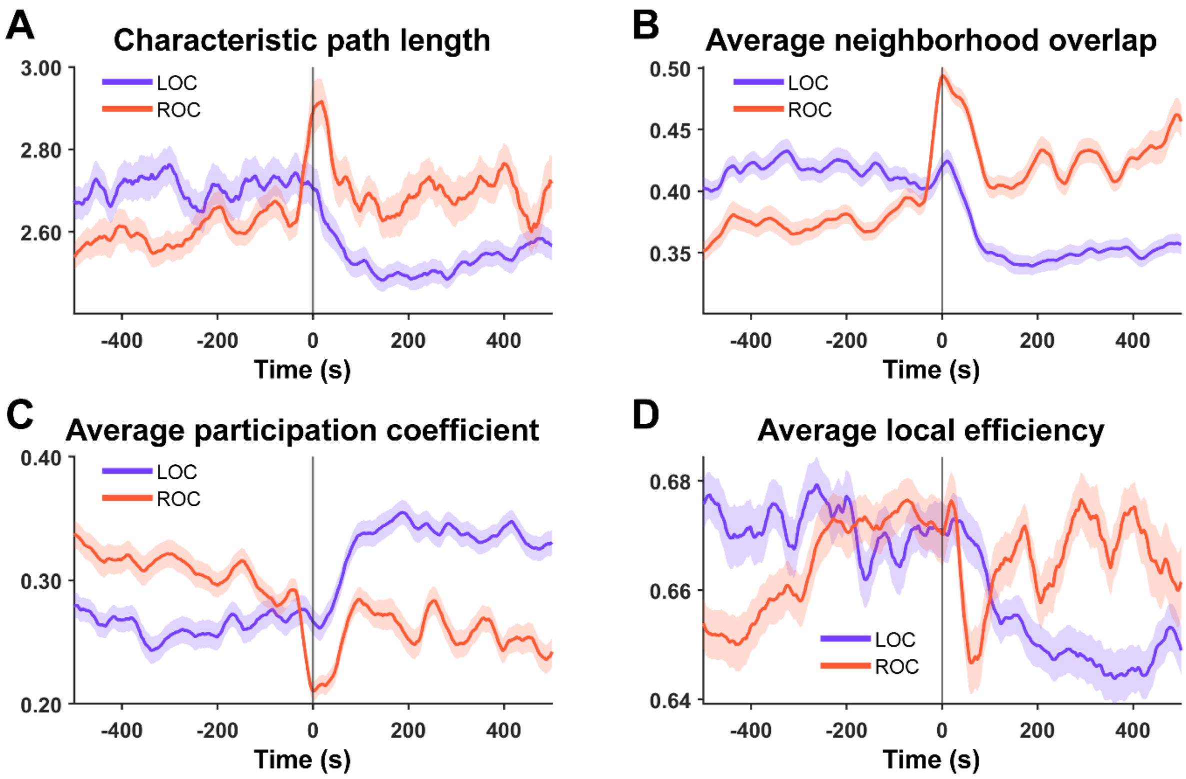


**Supplementary Figure 10. Global topological changes around LOC and ROC.**

Time courses of (**A**) characteristic path length, (**B**) average neighborhood overlap, (**C**) average participation coefficient, and (**D**) average local efficiency, aligned to pupil-defined LOC and ROC. Purple and orange lines indicate LOC and ROC, respectively; shaded regions show SEM across recording sessions. The vertical line marks the transition time.

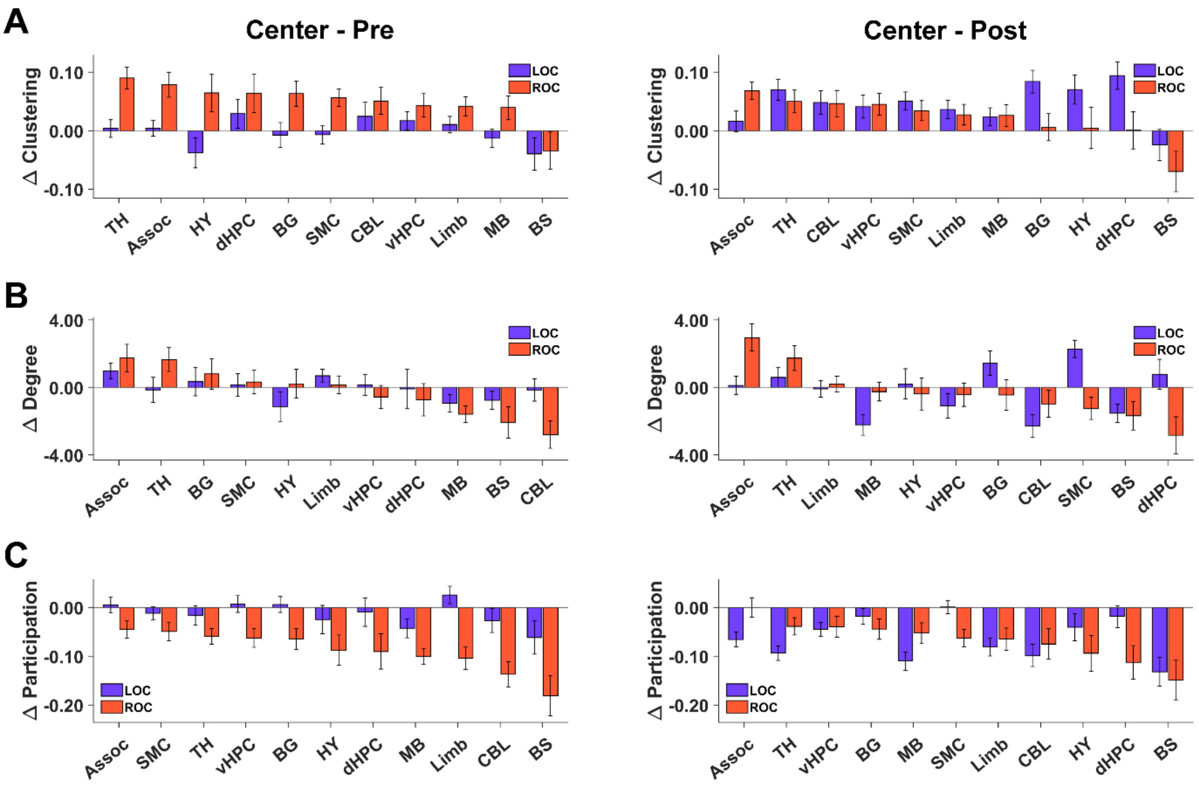


**Supplementary Figure 11. Transition-related nodal graph-metric changes across anatomical systems.**

System-level changes in nodal clustering coefficient (**A**), degree (**B**), and participation coefficient (**C**) for transition–pre (left) and transition–post (right). Bars and error bars show the mean and SEM across recording sessions.

**Captions for supplementary movies.**

Supplementary Movie 1: Whole-brain BOLD dynamics around LOC.

Supplementary Movie 2: Whole-brain BOLD dynamics around ROC.
